# Contextual dependencies expand the re-usability of genetic inverters

**DOI:** 10.1101/2020.07.15.204651

**Authors:** Huseyin Tas, Lewis Grozinger, Ruud Stoof, Victor de Lorenzo, Angel Goñi-Moreno

## Abstract

The design and implementation of Boolean logic functions in living cells has become a very active field within synthetic biology. By controlling networks of regulatory proteins, novel genetic circuits are engineered to generate predefined output responses. Although many current implementations focus solely on the genetic components of the circuit, the host context in which the circuit performs is crucial for its outcome. Here, we characterise 20 genetic NOT logic gates (inverters) in up to 7 bacterial-based contexts each, to finally generate 135 different functions. The contexts we focus on are particular combinations of four plasmid backbones and three hosts, two *Escherichia coli* and one *Pseudomonas putida* strains. Each NOT logic gate shows seven different logic behaviours, depending on the context. That is, gates can be reconfigured to fit response requirements by changing only contextual parameters. Computational analysis shows that this range of behaviours improves the compatibility between gates, because there are considerably more possibilities for combination than when considering a unique function per genetic construct. Finally, we address the issue of interoperability and portability by measuring, scoring, and comparing gate performance across contexts. Rather than being a limitation, we argue that the effect of the genetic background on synthetic constructs expand the scope of the functions that can be engineered in complex cellular environments, and advocate for considering context as a fundamental design parameter for synthetic biology.

## 1. INTRODUCTION

The abstraction of gene regulatory signals into on (high) and off (low) values allows for the design and implementation of genetic Boolean circuits^1^ inspired by digital electronics. Such devices result from assembling two or more genetic logic gates^1, 2^—the basic unit for processing information in genetic circuits based on Boolean logic. A core objective of Synthetic Biology^*3*^ is the building of new regulatory circuits to compute inputs into outputs according to predefined logical functions^*4*^, which are then used in a number of applications, ranging from bioproduction^*5*^ to medical diagnosis^*6*^. Although this approach has been relatively successful, genetic logic gates are way more fragile and less reliable than their electronic counterparts as their signals are rarely constant and often fluctuate over time^*7, 8*^. Consequently, the large-scale control of gene regulation based on Boolean logic alone is challenging. The central underlying issue is that a number of features intrinsic to biological systems, such as gene expression noise, analogue signalling^*9*^ and evolutionary dynamics^*10*^, make the intracellular environment an unsuitable domain for engineering idealised Boolean logic^*11*^.

A fundamental challenge for the design of robust synthetic circuits, which underpins this work, is the oversimplified model that DNA elements (i.e., gates) alone explains the performance of genetic circuits. Based on this assumption, the host chassis (the cell that receives a specific genetic construct) is generally ignored and the interplay of a genetic circuit with the host context is most often overlooked—an issue that has been identified essential for the predictability of synthetic biology devices^*12*^. Both the burden imposed by synthetic constructs on the host^*13, 14*^ and the impact of context on genetic activity^*15*^, have phenotypic implications that cannot be predicted from a gene-centric standpoint. Recently, the term host-awareness^*16, 17*^ has been coined to bring attention to this problem, which is at the core of the lack of part interoperability^*18*^ (i.e., parts that show similar performance in different host contexts).

While most synthetic biology efforts make use of only one host chassis to develop and characterise genetic constructs, potential applications may require the same genetic devices to work with different cell types^*19*^. For instance, circuit-constructs optimised in *Escherichia coli* for rapid prototyping, might be implanted into *Pseudomonas putida* for a bioremediation application^*20*^ or into *Geobacter sulfurreducens* for bioelectricity production^*21*^. However, circuit performance will likely differ in different chassis, gene dosages and vectors, highlighting the importance of context in host-circuit design. As a result, the performance of a given genetic logic device would not only be a consequence of its DNA sequence but also would be influenced by its context. Within this scenario, modifying the context could fine-tune the performance of logic gates, thus engineering reconfigurable genetic logic devices which share the same sequences but exhibit different behaviours^*22*^. In the work presented below we inspect these scenarios by analysing quantitatively the behaviour of a collection of genetic inverters in different strains of the same species, in other species and in either case with the same devices borne by low, medium and high-copy number vectors. The results expose that playing with these biological backgrounds expand the range of parameters that rule the behaviour of each construct. On this basis, we entertain that context-variability should be considered an advantage for circuit design rather than being seen as problematic.

## 2. RESULTS

### Generation of gate-context libraries

To generate enough data on the contextual dependencies of genetic inverters we made use of 20 NOT logic gates assembled with a suite of promoters and repressors first developed as components of the CELLO platform for *E. coli*^*1*^ and then recloned in broad host range vectors of different copy numbers for delivery to different types Gram-negative hosts^*23*^. The logic function (NOT or inverter) corresponds to a genetic device that expresses a target gene (output high) if it is not negatively regulated (input low) and vice-versa. The inverters used are pairs of a specific regulator (repressor) and its cognate promoter (Figure 1A). The characterised transfer functions measured the impact on promoter activity (output; captured by the expression level of an *ypf* reporter fused downstream) generated by specific concentration of regulator (input). In order to manipulate the expression level of the regulator, its coding sequence was placed under the control of a *lac* promoter, which was externally induced by IPTG. For gates characterisation, these were entered in a bacterial host, which was then used to measure the NOT function (Figure 1B). Both the IPTG concentration and the output *yfp* fluorescence were converted to relative promoter units (RPUs) to standardise the characterisations. The reference dataset of the behaviour of the 20 gates under inspection in *E. coli* NEB10β —12 main gates plus 8 variants—was retrieved from Nielsen et al^1^ (Table S1).

**Figure 1.**
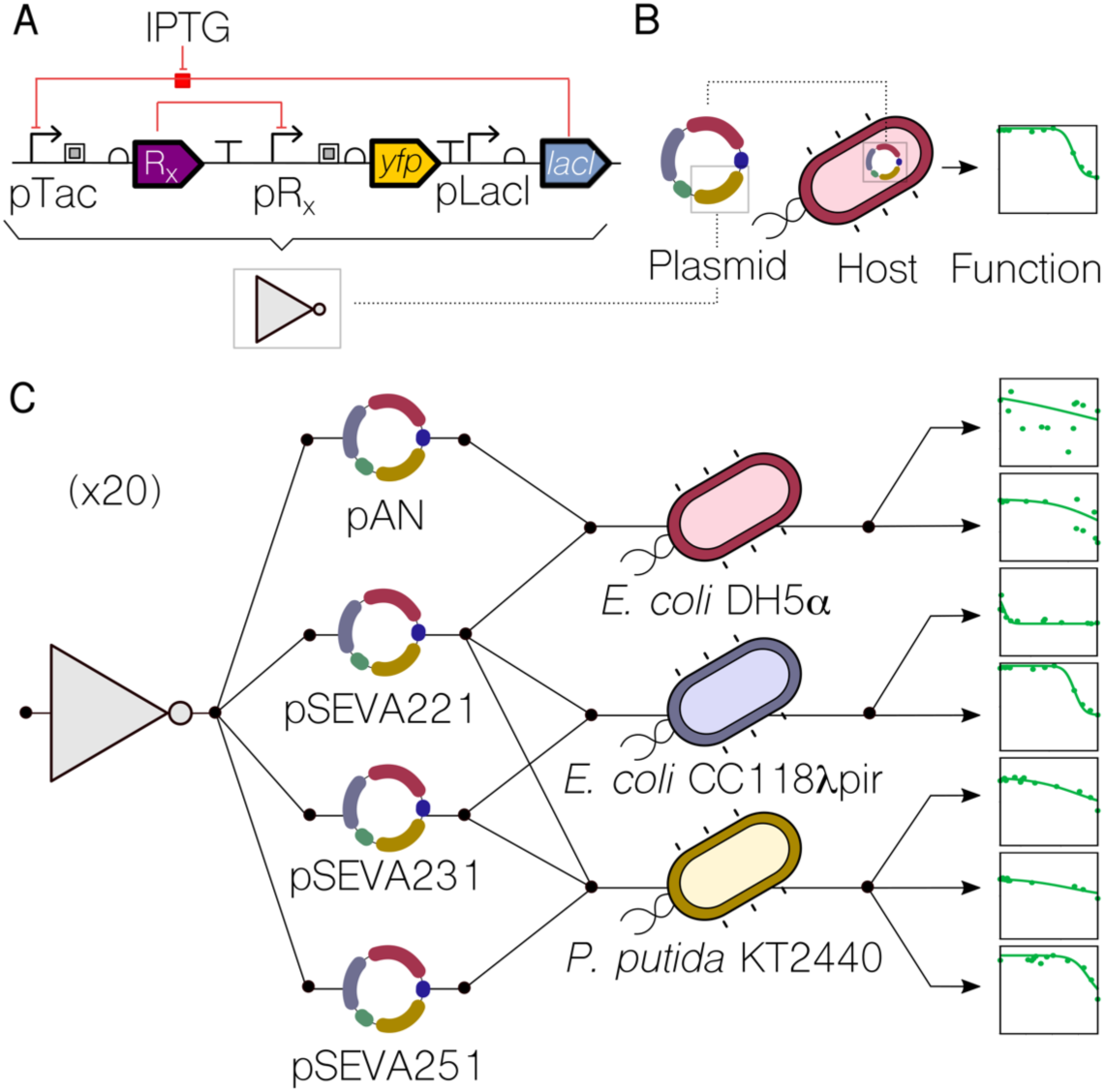
Generating a library of gate-context devices. **A**. Genetic inverters (NOT logic gates) were placed in between the pTac/LacI system (the input) and the *yfp* gene (the output). Key components are a repressor (R_x_) and its cognate promoter (pR_x_). **B**. For a genetic construct to be measured, it needs to be cloned into a plasmid which, in turn, is transformed into a host cell—thus using a single context. **C**. In this work, each genetic inverter (from an initial library of 20 gates) was measured in a number of context setups. These setups were based on combinations of 2 plasmid backbones (pAN and pSEVA), one of which with 3 different origins of replication—RK2 (221), pBBR1 (231), RFS1010 (251)—and 3 different hosts (*E. coli* DH5α, *E. coli* CC118λ*pir, P. putida* KT2440). The performance of the resulting 135 gate-context devices was characterized experimentally by using flow cytometry and analysed computationally to find the impact of contextual dependencies on inverter’s behaviour.

To assess the impact of the host context on gate performance, both the plasmid backbone and the cellular chassis were changed. As far as the carrier backbone is concerned, gates were cloned into the pAN and pSEVA^*24*^ backbones, considering different origins of replication that led to low (RK2, pSEVA221), medium (pBBR1, pSEVA231) and high (RFS1010, pSEVA251) copy numbers. This contextual feature accounted for dynamics generated by circuit burden^*25*^, since more copies of the same gate would increase the cost (of running it) to the cellular machinery. Regarding the chassis, we used two *Escherichia coli* (DH5α and CC118λ*pir*) strains and one *Pseudomonas putida* (KT2440) strain. Combinations of these resulted in a library of gate-backbone-host devices (Figure 1C) where the final performance cannot be explained by the genetics of the NOT logic gate alone. That is, the DNA sequence of the constructs is not enough to predict the behaviour of the gate—information about the context is then essential for understanding the genotype-to-phenotype dynamics. As shown in Figure 1C, each logic gate in this study can have up to 7 context-dependent dynamic behaviours, some of which differ significantly. Specifically plots shown in Figure 1C correspond to the characterisations of gate PhlF (one of the 20 gates of the initial library) in seven different contexts. While the performance changes abruptly in some cases (e.g., in contexts 3 and 4), it did not change significantly in others (e.g. in contexts 5 and 6), suggesting that contextual dependencies act as a hidden layer of parameters that must be carefully considered to achieve a predictable logic gate design—an issue which has been traditionally overlooked.

### Effects of cross-context portability

When a genetic logic gate is either passed onto another organism, or carried by a different backbone, the interplay between itself and the context changes^*26*^. Contextual dependencies are adjusted. These modifications alter the expression levels of a gate, its dynamic range and (in some cases) its logic function. Moreover, context-dependent changes of qualitative behaviour imply that the dynamics of the interplay between context and construct are nonlinear. That is, a given pair of gates may suffer similar modifications in one context but very different in another.

For example, PsrA-R1 and PhlF-P2 show these effects (Figure 2A). When both gates are hosted by chassis *E. coli* DH5α, the backbone (either pAN or pSEVA221) seems to play a key role in the logic outcome of PhlF-P2, which becomes more step-like with pSEVA221 (i.e. sharper transition from on to off). In contrast to this, gate PsrA-R1 does not follow that trend and remains qualitatively unchanged, although absolute expression values drop. Using the same backbone (pSEVA221) we then tested the context impact of varying the *E. coli* strain. Whilst the performance of PhlF-P2 is qualitatively the same (with smaller dynamic range), PsrA-R1 shows a qualitative change, becoming more step-like, thus showing more desirable behaviour than in other contexts. These inconsistencies in changes of qualitative behaviour of gates highlight the difficulty of compensating for such effects in order to engineer context-independent circuits^*26*^. However, there are also more predictable contextual changes in which that strategy may work well. For example, when both gates are hosted by *E. coli* CC118λ*pir*, changing the backbone from pSEVA221 to pSEVA231 (that only differ in the origin of replication) generates almost the exact same phenotypic modification. Finally, a marked difference occurs when the gates are hosted by *P. putida* KT2240. In these contexts, the gates lose their NOT logic, regardless of choice of backbone (pSEVA221, pSEVA231 and pSEVA251). The characterisation of the full library (20 gates) is shown in Supplementary Information Table S2.

**Figure 2.**
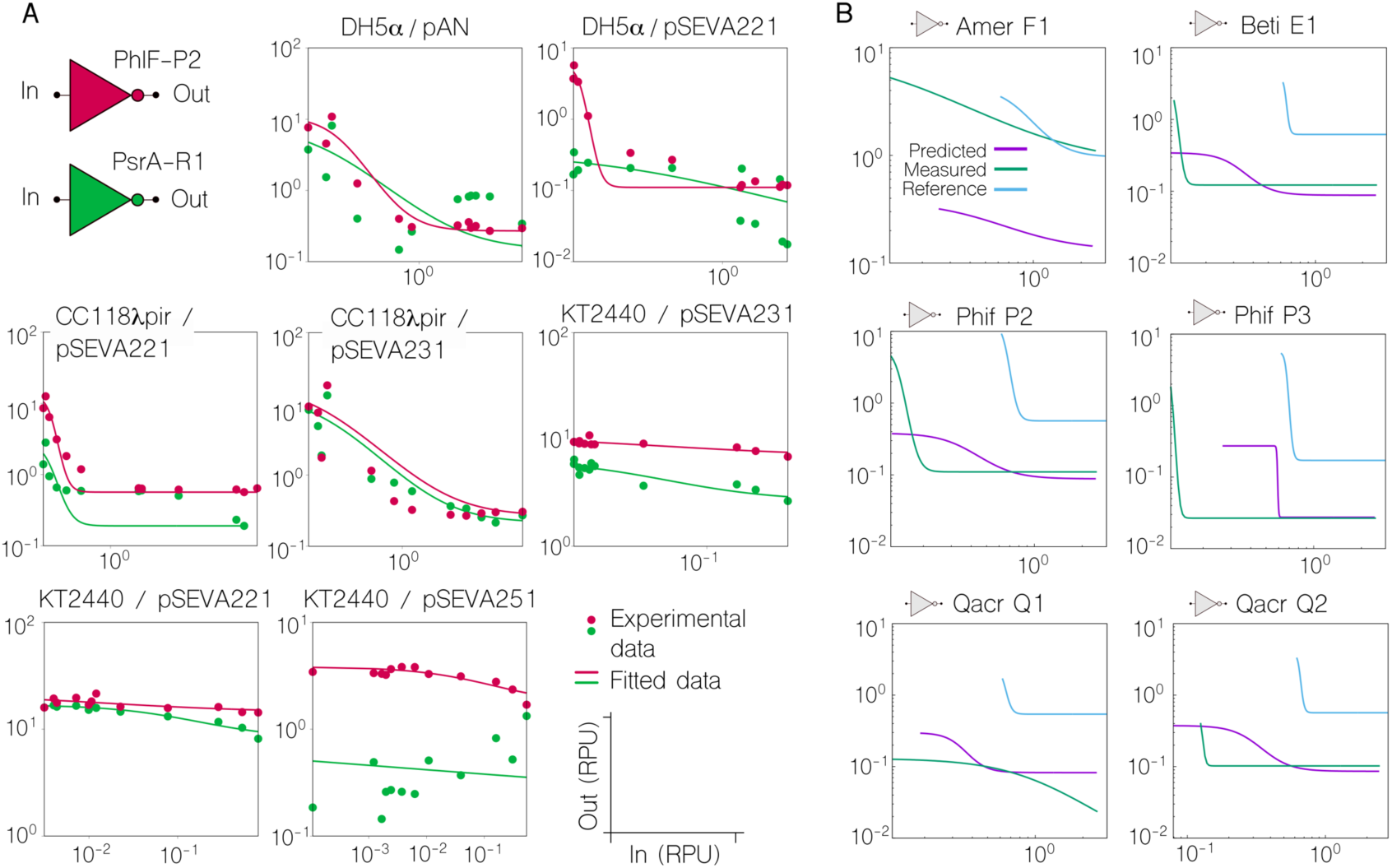
Non-linear effects in the cross-context portability of inverters. **A**. Plots comparing the characterisation of two gates, PsrA-R1 and PhIF-P2, in different contexts. As well as each individual characterisation is differing across contexts, the relationship between the two characterisations also differs, depending on both strain and plasmid. This is, some contextual changes impact on a similar way on the performance of two inverters, while others impact on a different way—what we refer to as non-linear modifications. **B**. Non-linearities made the prediction of gate performance changes between contexts an overarching challenge. Predictions were made for gates in the *E. coli* DH5α (pSEVA221) context (‘Predicted’ line), based on their characterisations in *E. coli* CC118λ*pir* (pSEVA221) i.e. ‘Reference’ line. The actual characterisation of the gate is shown for comparison (‘Measured’ line). The gate upon which these predictions are based is AmtR-A1. It can be seen that the linear transformation used for prediction cannot accurately produce all the differences in behaviour seen between contexts. For example, although translations in the Input and Output axis appear to be predicted well in some cases (see for example QacR-Q2), more qualitative changes in the shape of the response curve cannot be addressed by this linear transformation (see for example QacR-Q1).

As a result of this non-linear performance in cross-context experiments, the issue of inter-context predictions arose as a formidable challenge. For example, an attempt to predict the performance that a number of gates would display into the context *E. coli* DH5α (pSEVA221)—host and plasmid—could not match all experimental data, revealing differences where the non-linear patters were stronger (Figure 2B). For this prediction, we took as a reference the performance of the same gates into the context *E. coli* CC118λ*pir* (pSEVA221) i.e., only change in host cell, and the measurement of only one of the gates (AmtR-A1) into the new context *E. coli* DH5α (pSEVA221). With the information provided by the characterisation of AmtR-A1 in both context setups, we applied linear transformations to predict the performance of the rest of the gates into the new scenario. As expected, some of the gates showed a relatively good prediction (good candidates for portability applications), but that was not the case for all of the constructs. Although predictable gate portability is then highlighted as an open problem, contextual dependencies offer a unique opportunity for fine-tuning gate performance, which we carefully analysed as explained next.

### Enhanced gate compatibility by fine-tuning contextual dependencies

Building a genetic circuit by coupling genetic logic gates requires an assessment of their compatibility, to determine which gates can be connected. In order to connect two gates, the output levels of one must match the input levels for the other. If not, it may result in failure of the overall circuit logic^*1, 27, 28*^. This is one of the major bottlenecks that restrict the depth of genetic logic circuits and limit scalability, since not every gate within a library will be compatible with another. The analysis of inter-gate compatibility is therefore fundamental for circuit design and is an integral part of current synthetic biology Computer Aided Design tools^*1, 29*^. However, knowledge about the effect of context on gate compatibility has until now been lacking.

In order to tackle this contextual issue, we first scored the matching of all the gate pairs in the library according to their input and output thresholds (Figure 3A). The inclusion of the input thresholds in the output ones defines a pair as “compatible”; each pair of gates in the library was categorised as such (see Methods for details of this calculation). Moreover, the information provided by the compatibility was complemented by the introduction of a similarity score (Figure 3B). While the former relates two different gates, the latter relates the same gate to itself when varying contextual dependencies. This score quantifies the impact of specific context variations on each gate.

**Figure 3.**
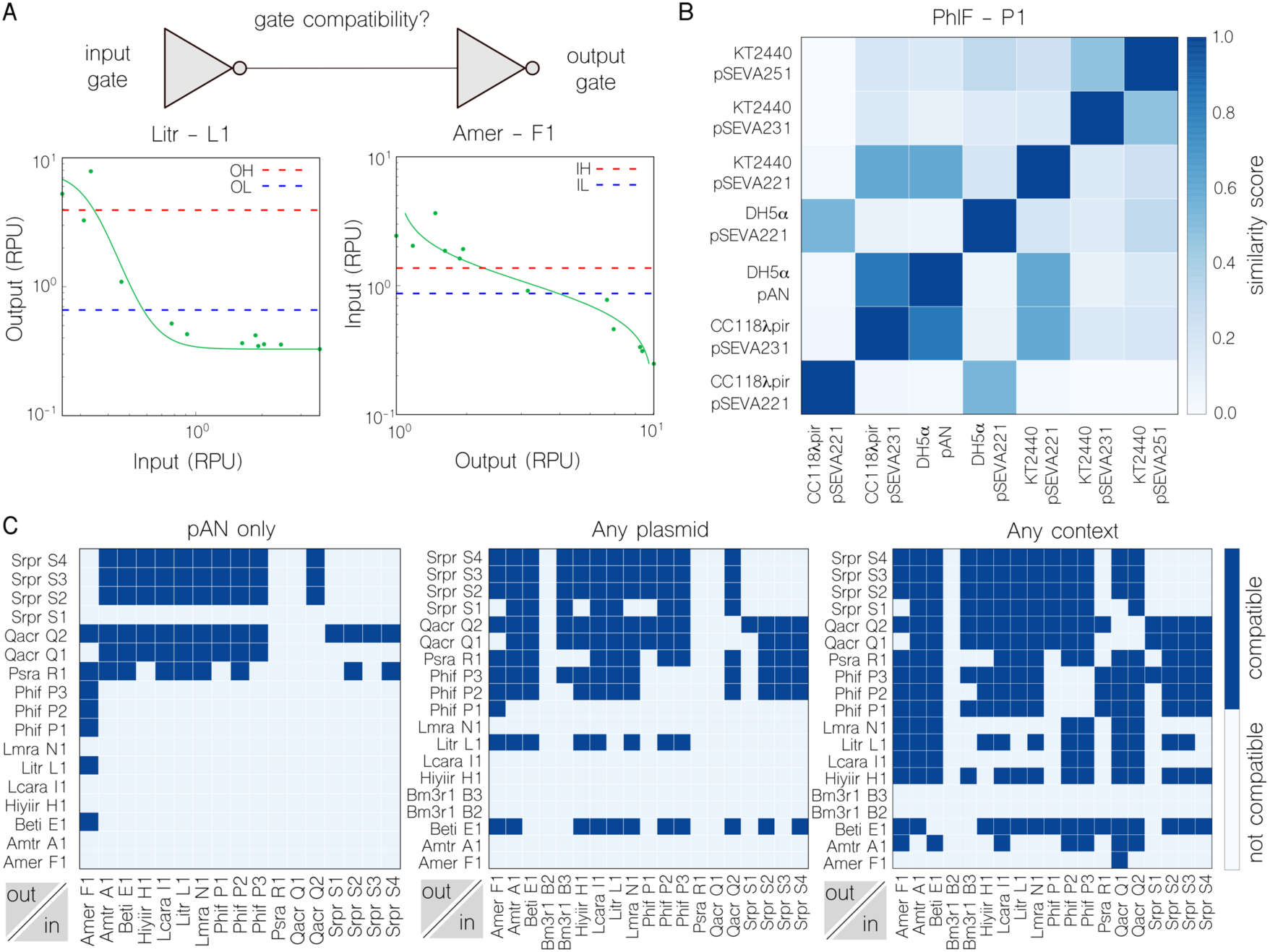
Comparing inverter compatibility and similarity across contexts. **A**. Gate compatibility indicates if two gates can be sequentially assembled—the output of the first gate is compatible with the input of the second—or not. Since the IH (Input High) and IL (Input Low) thresholds of the output gate, AmeR-F1, lie between the OH (Output High) and OL (Output Low) thresholds of the input gate, LitR-L1, this pairing is compatible. **B**. A heatmap of similarity scores (which refers to how similar the shape of both inverter’s transfer function is) calculated using discrete Frechet distance between the characterisation of PhIF-P1 in each of the seven contexts (darker is more similar). Most values within the score scale are covered, which highlights context contribution to final gate behaviour. **C**. Maps of compatible pairs for the gates characterised in: the strain *E. coli* DH5α with pAN as the only plasmid for all inverters (left), the strain *E. coli* DH5α with any variation in plasmid type (middle) and in any context choice (right). Considerably more compatible pairs are found when freedom is given in the choice of backbone, rising from 67 (left) to 203 (middle) pairs. The freedom to use both backbone and strain (right) as a design parameter yields the most compatible pairs (697) and maximum utilisation of gate combinations in the library (∼68%).

In this analysis, constructs were considered as gate-context entities (e.g. *E. coli* DH5α (pAN::PsrA-R1) or *E. coli* DH5α (pSEVA221::PsrA-R1), rather than individual gates alone (e.g. PsrA-R1) so that results account for the performance of a gate in a given context. We consider that high numbers of compatible pairs in a library is a desirable trait, and examine the impact of the two contextual features we focus on (backbone and host) on this metric, both independently and together. Figure 3C (left) shows the results of the former, where the compatibility is assessed in the case of gates sharing backbone pAN and host *E. coli* DH5α. In this scenario, the number of compatible pairs found within the library was 67. However, when we allowed the calculation algorithm to consider all possible backbone options (while keeping strain constant), the number of compatible pairs increases to 203 (a 303% increase, Figure 3C middle). We conclude that consideration of backbone as a design parameter results in a more flexible, and reconfigurable, library with the ability to include dynamics that are not captured by just DNA sequences e.g., the copy number of circuits (thus their burden to the cell).

Gate compatibility was enhanced further by allowing the algorithm to consider the host as a design parameter, as well as backbones (Figure 3C right). Although the number of compatible pairs relative to the total number of pairings being evaluated remained almost constant at roughly 2%, the absolute number increased to 697. Due to the consideration of contextual dependencies, the original gate library of only 20 genetic devices was increased to 135 different functions. Therefore, there were many more options to evaluate and more compatible pairs found. However, some of these pairs correspond to gates that are compatible only if they are inside different hosts. For example, the gates HIyIIR-H1 and AmeR-F1 can only be matched (i.e., their function is complementary) if the former is hosted by *E. coli* DH5α and the latter by *E. coli* CC118λ*pir*. This suggests that taking a multicellular (distributed) computing approach^*30-33*^ may be a suitable strategy if coupling the functions of these two constructs. In multicellular computations, a predefined function is distributed across different engineered bacterial strains (or species), which are connected in such a way that the output of one cell is the input of another one. Therefore, considering the host of a genetic construct within circuit design will allow for building both intra-and inter-cellular computations^*11*^.

### Context-aware design rules for layered logic gates

The design of synthetic genetic circuits typically overlooks contextual features by considering that phenotypic performance can be explained by the DNA sequence of the synthetic construct alone. However, this over-simplification has negative implications; for example, it requires considerable effort to adapt a genetic circuit to a new host^*34*^. The fact that genetic constructs show different dynamics depending on their context is not necessarily a disadvantage for pre-defined circuit design—could we rationally use such variability? To begin to address this question, we carried out computations in order to identify the circuit depth (i.e., number of layers^*35*^) that could be achieved by connecting gates within our library, and assessed the impact of contextual effects in such a chain (Figure 4). First, when considering all gates in the same context, with backbone pSEVA221 and hosted by *E. coli* CC118λ*pir*, the maximum depth was 3 (Figure 4A). That is, there are 3 gates that can be connected consecutively while maintaining the correct logic output (i.e., logic values 0/1 are effectively transmitted from beginning to end). Every other valid configuration will result in fewer (or the same) number of layers. We find that increasing the number of contexts available can significantly increase the maximum depth computed by the search algorithm. As shown in Figure 4B, allowing another context by including gates characterised with any backbone (but still hosted by *E. coli* CC118λ*pir*) increases the maximum depth to 5. This can be further improved upon by allowing freedom in the choice of host, for a total of 7 contexts (Figure 4C). In this case, the computed maximum depth is 11 — a circuit depth that is far beyond the current state-of-the-art for synthetic circuitry^1^.

**Figure 4.**
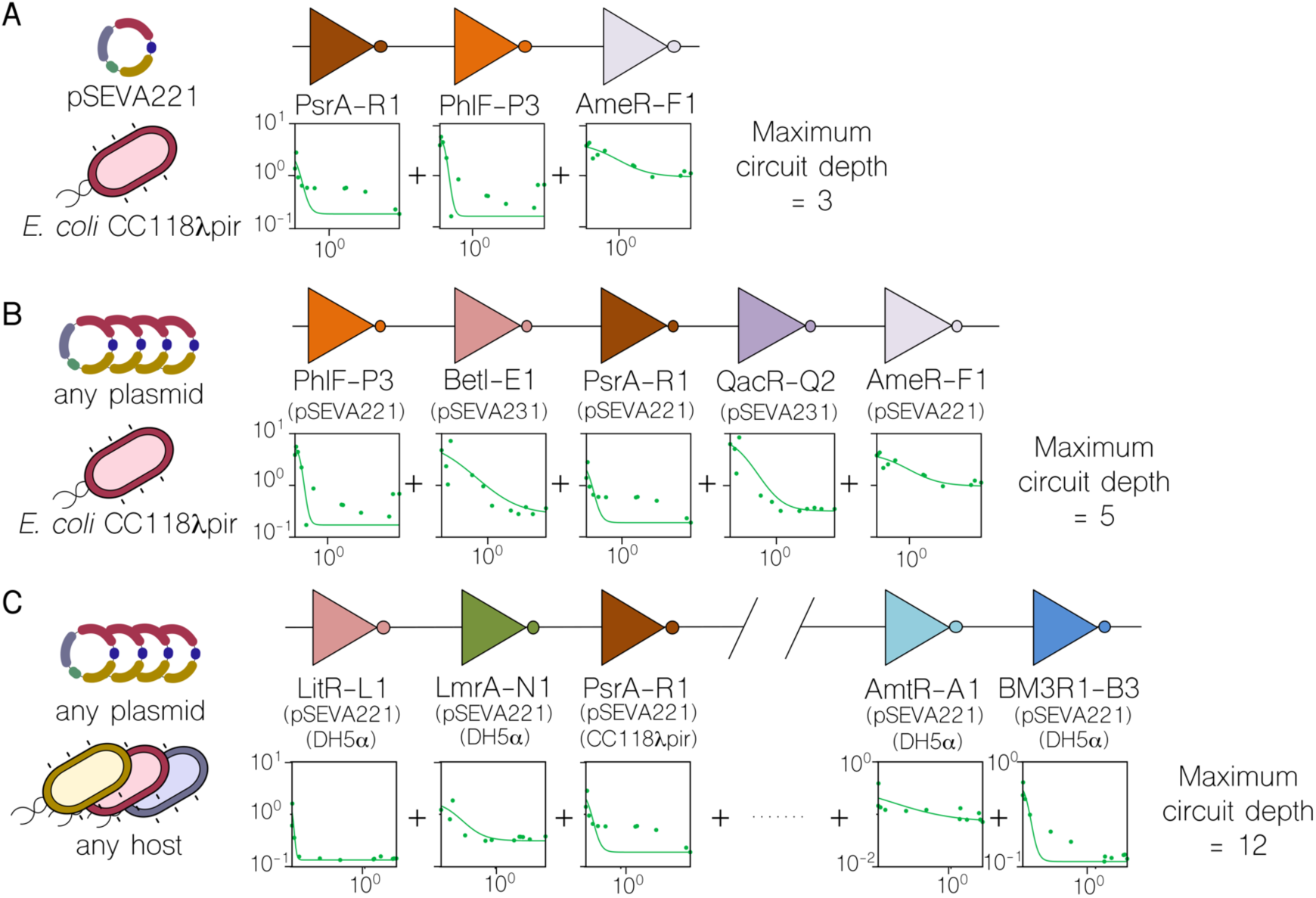
Calculation of maximum circuit depth as a result of layering inverters. Based on the compatibility between gates, these were layered within the library in order to evaluate the impact of contextual dependencies on circuit size. **A**. The maximum depth calculated when the computational method is forced to consider all gates carried by the low copy-number plasmid pSEVA221 and hosted by *Escherichia coli* CC118λ*pir*, is 3 gates-deep. **B**. If the algorithm is free to select any plasmid (but still forced to *E. coli* CC118λ*pir*), the maximum depth increases to 5. In this scenario, two gates are carried by the medium copy-number plasmid pSEVA231. **C**. In the last analysis, the calculation used all contextual dependencies, including the variation in host chassis. The maximum number of gates layered increases to 12 (only 5 shown in figure—refer to Supplementary Material for more information). In the sketch shown in the figure, 4 out of the 5 gates were characterised in the strain *Escherichia coli* DH5α. For all graphs: x-axis refers to the input and y-axis to the output (both RPU).

## DISCUSSION

A fundamental driving force for synthetic biology^*3, 36*^ is the clarification of mechanistic assumptions as our understanding of molecular processes increases, which allows scientists to add novel tools to the catalogue for engineering living systems. Although the cellular environment consists of much more than DNA, circuit design^*1, 29*^ typically revolves around genetic elements (promoters, terminators, RBSs…) in order to link genotype to phenotype—an over-simplified reductionist approach. The comfortable, yet error-prone, assumption that engineered parts alone can ultimately explain phenotypic performance needs to be expanded upon^*12*^. This leads us to consider what has been termed genetic background^*15*^ and host-aware^*16*^ dynamics: cellular features and constraints that have an impact on circuit performance but are not captured by the DNA sequences of the construct. In recent years, several of these features have been analysed: the impact of having limited cellular resources^*37, 38*^ (e.g. ribosomes) to “spend” on synthetic constructs, the effects of placing DNA parts in different genomic locations^*39, 40*^, the role played by metabolism in genetic control^*41, 42*^, or even genetic stability^*43*^ due to evolution over time. All these effects turn the portability of genetic circuits into an overarching challenge—the fine-tuning of a circuit to work inside a different host (to the one it was originally built-in) is still a major task^*26*^. Furthermore, it limits the scope of biological circuits by solely using a DNA-insert toolbox for designing circuits.

Here, we use the word “context” to refer to the molecular background of the cell beyond genes and analyse how such context can be used for improving biocircuit design. By “dependencies”, we mean the constraints imposed by the context on a given genetic construct. Therefore, genetic logic gates are exposed to contextual dependencies that influence their phenotypic behaviour. Although synthetic biology is a field full of metaphors^*44*^ already, we entertain here a new one that we consider to provide a useful conceptual frame: the use of contextual dependencies as in a software engineering problem. Any piece of software, or program, must run inside a specific environment (e.g. operating system) and software engineers usually face the problem of adapting it to the particular dependencies of the environment/context at stake. Under this metaphor, genetic circuits are considered software (instead of hardware^*45*^) whose performance is deeply linked to context-specific dependencies, which can allow designers to access functions that could not be coded otherwise. In this paper, we propose that contextual dependencies are important parameters for circuit design, and focus on [i] backbone carrying the construct, and [ii] cellular host in which the construct performs.

In this work we exploited a library of 20 genetic inverters (NOT logic gates), which are combined with 4 different backbones and 3 cellular strains to give a total of 135 gate-context constructs. In this regard, the number of functions exposed by the library increases by 675% due to the addition of these two contextual dependencies. With this new library we carried out experiments in order to assess the implications of adding context to the context-free initial collection of NOT gates. First, the characterisation of the constructs showed how gate behaviour changed across contexts in a nonlinear fashion. That is, the phenotypic modifications in the performance of one gate across two contexts may not match those of another gate under the same contextual transformations. This has major implications for the portability of genetic devices, since not all genetic components may be affected in the same way upon host change—thus building complex portable devices will become difficult (if not entirely impossible). Second, our experiments suggested that the compatibility of gates (so that they could be composed; the output of the first being the input of the second) does not only depend on selected genetic inserts, but also on their context. While only 67 compatible pairs were found in the original library of 20 inverters, the number increased to 697 in the new library. For instance, by allowing gates to be carried by 4 different backbones, the computational algorithm was able to evaluate the compatibility of 4 functions instead of 1 and return not only the name of the compatible gate but also the name of the backbone to use for carrying it. This allows reconfiguration of genetic constructs, since the same piece of DNA-insert can have different behaviours depending on rationally selected contextual dependencies. Finally, the use of the cellular host as a separate design parameter allowed identification of gate pairs that were only compatible if connected gates were located in different strains/species. This suggests that multicellular computing approaches^*30, 46, 47*^ may be best suited for a given set of functions, and establishes rational criteria for the selection of cellular chassis in such distributed consortia from a bottom-up design.

In a similar way to living systems that use a number of mechanisms to go from genotype to phenotype, we advocate for the development of genetic circuits by considering whole-cell dynamics—including contextual dependencies. This will result in the design of biological circuits that are closer to the internal workings of natural systems^*11*^—therefore more robust, reliable, predictable and reproducible.

## METHODS

### DNA and Strain Construction

All cloning steps were done in *E. coli* CC118λ*pir*. Primers are ordered from Merck Sigma Aldrich, Inc. The repressible and inducible systems were previously described by Voigt Lab and acquired by the courtesy of Voigt Laboratory in MIT (USA). Description of the 20 different NOT gates moved into broad host range pSEVA backbones are described in Tas et al.^*23*^. Components of the original inverters, like terminators, RBSs, insulators, etc. were kept the same during the SEVA conversion. SEVA backbones have two terminators, T0 and T1 which are important to lower potential leakages. Required oligo list can be found in the Supplementary Information (Table S3). The pAN backbone^*1*^ has a kanamycin resistance gene with a p15A origin of replication which is ∼15 copy number in *E. coli* NEB10β strain.

### Medium and Experimental Protocols

In all experiments (unless stated otherwise) M9 minimal medium for *E. coli* and M9 medium for *P. putida* were used. The ingredients of the M9 medium used are as following: for 250 ml of liquid medium, 25 ml 10X M9 salts, 500 µl of 1M MgSO_4_, 2.2 ml of 20% carbon source (glucose for E. coli and citrate for P. putida), 125 µl of 1% Thiamine, 2.5 ml of 1% Casamino acids and milliQ-H_2_O up to 250 ml. Concentration of kanamycin used is 50 *µ*g ml^*–*1^ in the experimentation procedures. IPTG was used as inducer for pTac/LacI inducible system in 12 different concentrations diluted from 1M stock concentration that are 0, 5, 10, 20, 30, 40, 50, 70, 100, 150, 200, 500 and 1000 µM. For synchronizing the cells in the experimental procedure, cultures are started from a single colony picked from LB agar plate which is each time freshly prepared from −80C glycerol stock by inoculating it O/N in 1 ml M9 minimal medium. O/N cultures after saturation were diluted by ∼666 times to inoculate 200 µl M9 minimal medium in 96 well plate for 24h, which is enough to reach to 0.2 - 0.3 OD in 96 well plate after which for halting the growth cells were kept on cold platform during the measurements.

### Flow Cytometry Analysis

Miltenyi Biotec MACS flow cytometer at channel B1 with an excitation of 488 nm and emission of 525/50 nm was used for measuring YFP fluorescence distribution of each sample. 30000 events were defined as the statistically sufficient amount under singlet gating for each sample. Calibration of the flow cytometer was done daily by using MACSQuant Calibration Beads. Throughout flow cytometer measurements samples were always kept on cold 96 well plate platforms. For the analysis of the data FlowJo software was used. In the analysis, gating was done via usage of auto-option and allowing to cover at least 50% of the whole events run while Forward and Side scatters were plotted, and the same gating conditions were kept for all samples in the same group.

### Fluorescence data pre-filtered by cell size

In order to unify fluorescence measures between and within flow cytometry experiments, we analysed fluorescence and scattering values. Variation in cell size across experiments showed that median fluorescence values were decisively affected, therefore inaccurate for the sake of comparison. To compare between experiments, we took the distribution of fluorescence for single scattering values. A full description of this process is detailed in Supplementary Information.

### Standard fluorescence measurements

Two extra plasmids were used for measurements, the autofluorescence plasmid (Backbone::1201), and the reference standard plasmid (Backbone::1717) that triggers *yfp* expression under the pLacI constitutive promoter. In order to derive reference promoter units (RPU), the following equation was applied:

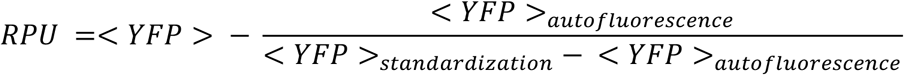

<YPF> stands for the median fluorescence value of the inverter that is to be standardised into RPU, <YPF>_autofluorescence_ is the median fluorescence value of the auto-fluorescence plasmid, <YPF>_standardisation_ indicates the median fluorescence value from the standardisation plasmid. RPU values were calculated in transfer function plots for different data points using at least 6 inducer levels covering the range of induction up to saturation.

### Data fitting

The pre-filtered experimental data were fitted to a 4-parameter hill equation of the form:

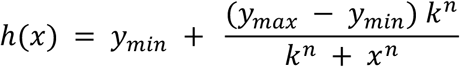

The parameter values for y_min_ and y_max_ were set to the minimum and maximum of the corrected experimental data. The values for k and n were then fitted using the least squares method from the scipy.optimize Python package^*48*^, with logarithmic residuals.

### Calculating compatibility

Thresholds OL, OH, IL and IH were computed from the parameters of the fitted hill curves according to the definitions given in [1]. OL and OH are twice y_min_ and half of y_max_, respectively. IL and IH are the values of x for which the output of the fitted hill function is equal to OL and OH, respectively. Accordingly, the values of IL and IH were calculated with the following formulae:

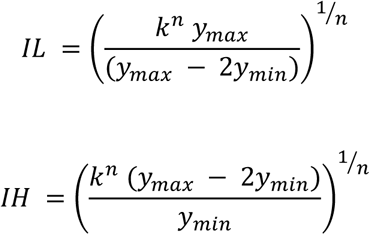

Inverters were considered operational under the condition that OH>OL and IH>IL for their fitted hill curve. For a pair of operational inverters, A and B, their compatibility score was defined as:

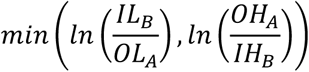

with the implication that inverter A can be connected as input to inverter B, if and only if their compatibility score is positive.

### Computation of inverter chains

Chains of compatible inverters were found by creating a table of compatibility between available inverters, for which the entry for a compatible pair was 1, and all other entries were 0. This table was then treated as the adjacency matrix of the graph of all possible connections, and the longest paths were enumerated using a depth-first search of the graph. Paths in which the same repressor was used more than once were excluded from the results, thus imposing an upper bound of 12 on path length.

### Similarity measure

The discrete Frechet distance^*49*^ was used to measure similarity of the shapes of two experimental curves, after first log transforming and min-max normalisation of the data along both axes. The Frechet distance was then subtracted from 1 in order to produce a metric that increases as the shape of the curves becomes more similar. The discrete Frechet distance was computed using the ‘similarity measures’ Python package ^*50*^.

### Prediction

The goal of the prediction is to transform the characterisation of gates in a source context, to a characterisation in the target context. A single operable gate was selected arbitrarily upon which to base the prediction. The ‘scipy.optimize’ Python package^*48*^ is used to compute a linear transformation matrix, which when applied to the source characterisation, minimises the L1Loss between the transformed characterisation and the target’s true characterisation. Predictions for other gates in the library are then made by applying the same transformation to their characterisation in the source context.

## ACKNOWLEDGMENTS

Authors ate indebted to Esteban Martinez-Garcia for his feedback on this Project. This work was funded by the *SETH (*RTI2018-095584-B-C42) *(MINECO/FEDER)*, SYCOLIM (PCI2019-111859-2 ERA-COBIOTECH 2018) *Project of the Spanish Ministry of Science and Innovation*. MADONNA (H2020-FET-OPEN-RIA-2017-1-766975), BioRoboost (H2020-NMBP-BIO-CSA-2018/ 820699), SYNBIO4FLAV (H2020-NMBP/0500) and MIX-UP (H2020-Grant 870294) Contracts of the European Union, the InGEMICS-CM (S2017/BMD-3691) Project of the Comunidad de Madrid - European Structural and Investment Funds - (FSE, FECER), and the SynBio3D project of the UK Engineering and Physical Sciences Research Council (EP/R019002/1).

## REFERENCES

[1] Nielsen, A. A., Der, B. S., Shin, J., Vaidyanathan, P., Paralanov, V., Strychalski, E. A., Ross, D., Densmore, D., and Voigt, C. A. (2016) Genetic circuit design automation, Science 352, aac7341.

[2] Wang, B., Kitney, R. I., Joly, N., and Buck, M. (2011) Engineering modular and orthogonal genetic logic gates for robust digital-like synthetic biology, Nature communications 2, 1–9.

[3] Ausländer, S., Ausländer, D., and Fussenegger, M. (2017) Synthetic biology—the synthesis of biology, Angewandte Chemie International Edition 56, 6396–6419.

[4] Amos, M., and Goñi-Moreno, A. (2018) Cellular computing and synthetic biology, In Computational matter, pp 93–110, Springer.

[5] de Lorenzo, V., Prather, K. L., Chen, G. Q., O’Day, E., von Kameke, C., Oyarzún, D. A., Hosta-Rigau, L., Alsafar, H., Cao, C., and Ji, W. (2018) The power of synthetic biology for bioproduction, remediation and pollution control, EMBO reports 19.

[6] Slomovic, S., Pardee, K., and Collins, J. J. (2015) Synthetic biology devices for in vitro and in vivo diagnostics, Proceedings of the National Academy of Sciences 112, 14429–14435.

[7] Kaern, M., Elston, T. C., Blake, W. J., and Collins, J. J. (2005) Stochasticity in gene expression: from theories to phenotypes, Nature Reviews Genetics 6, 451–464.

[8] Bornholdt, S. (2008) Boolean network models of cellular regulation: prospects and limitations, Journal of the Royal Society Interface 5, S85–S94.

[9] Daniel, R., Rubens, J. R., Sarpeshkar, R., and Lu, T. K. (2013) Synthetic analog computation in living cells, Nature 497, 619–623.

[10] Sleight, S. C., Bartley, B. A., Lieviant, J. A., and Sauro, H. M. (2010) Designing and engineering evolutionary robust genetic circuits, Journal of biological engineering 4, 12.

[11] Grozinger, L., Amos, M., Gorochowski, T. E., Carbonell, P., Oyarzún, D. A., Stoof, R., Fellermann, H., Zuliani, P., Tas, H., and Goñi-Moreno, A. (2019) Pathways to cellular supremacy in biocomputing, Nature communications 10, 1–11.

[12] Cardinale, S., and Arkin, A. P. (2012) Contextualizing context for synthetic biology–identifying causes of failure of synthetic biological systems, Biotechnology journal 7, 856–866.

[13] Ceroni, F., Algar, R., Stan, G.-B., and Ellis, T. (2015) Quantifying cellular capacity identifies gene expression designs with reduced burden, Nature methods 12, 415.

[14] Gyorgy, A., Jiménez, J. I., Yazbek, J., Huang, H.-H., Chung, H., Weiss, R., and Del Vecchio, D. (2015) Isocost lines describe the cellular economy of genetic circuits, Biophysical journal 109, 639–646.

[15] Phillips, K. N., Widmann, S., Lai, H.-Y., Nguyen, J., Ray, J. C. J., Balázsi, G., and Cooper, T. F. (2019) Diversity in lac Operon Regulation among Diverse Escherichia coli Isolates Depends on the Broader Genetic Background but Is Not Explained by Genetic Relatedness, mBio 10.

[16] Boo, A., Ellis, T., and Stan, G.-B. (2019) Host-aware synthetic biology, Current Opinion in Systems Biology.

[17] Darlington, A., and Bates, D. G. (2016) Host-aware modelling of a synthetic genetic oscillator, In 2016 38th Annual International Conference of the IEEE Engineering in Medicine and Biology Society (EMBC), pp 1463–1466, IEEE.

[18] Pray, L., Relman, D. A., and Choffnes, E. R. (2011) The science and applications of synthetic and systems biology: workshop summary, National Academies Press.

[19] Kim, J., Salvador, M., Saunders, E., González, J., Avignone-Rossa, C., and Jiménez, J. I. (2016) Properties of alternative microbial hosts used in synthetic biology: towards the design of a modular chassis, Essays in biochemistry 60, 303–313.

[20] Dvořák, P., Nikel, P. I., Damborský, J., and de Lorenzo, V. (2017) Bioremediation 3.0: engineering pollutant-removing bacteria in the times of systemic biology, Biotechnology advances 35, 845–866.

[21] Ueki, T., Nevin, K. P., Woodard, T. L., and Lovley, D. R. (2016) Genetic switches and related tools for controlling gene expression and electrical outputs of Geobacter sulfurreducens, Journal of industrial microbiology & biotechnology 43, 1561–1575.

[22] Goñi-Moreno, A., and Amos, M. (2012) A reconfigurable NAND/NOR genetic logic gate, BMC systems biology 6, 126.

[23] Tas, H., Goni-Moreno, A., and de Lorenzo, V. (2020) A standardized broad host range inverter package for genetic circuitry design in Gram-negative bacteria, bioRxiv, 2020.2007.2014.202754.

[24] Martínez-García, E., Goñi-Moreno, A., Bartley, B., McLaughlin, J., Sánchez-Sampedro, L., Pascual del Pozo, H., Prieto Hernández, C., Marletta, A. S., De Lucrezia, D., and Sánchez-Fernández, G. (2020) SEVA 3.0: an update of the Standard European Vector Architecture for enabling portability of genetic constructs among diverse bacterial hosts, Nucleic acids research 48, D1164–D1170.

[25] Wu, G., Yan, Q., Jones, J. A., Tang, Y. J., Fong, S. S., and Koffas, M. A. (2016) Metabolic burden: cornerstones in synthetic biology and metabolic engineering applications, Trends in biotechnology 34, 652–664.

[26] Kushwaha, M., and Salis, H. M. (2015) A portable expression resource for engineering cross-species genetic circuits and pathways, Nature communications 6, 1–11.

[27] Prindle, A., Selimkhanov, J., Li, H., Razinkov, I., Tsimring, L. S., and Hasty, J. (2014) Rapid and tunable post-translational coupling of genetic circuits, Nature 508, 387–391.

[28] Gorochowski, T. E., Espah Borujeni, A., Park, Y., Nielsen, A. A., Zhang, J., Der, B. S., Gordon, D. B., and Voigt, C. A. (2017) Genetic circuit characterization and debugging using RNA-seq, Molecular systems biology 13, 952.

[29] Appleton, E., Madsen, C., Roehner, N., and Densmore, D. (2017) Design automation in synthetic biology, Cold Spring Harbor perspectives in biology 9, a023978.

[30] Regot, S., Macia, J., Conde, N., Furukawa, K., Kjellén, J., Peeters, T., Hohmann, S., De Nadal, E., Posas, F., and Solé, R. (2011) Distributed biological computation with multicellular engineered networks, Nature 469, 207–211.

[31] Goñi-Moreno, A., Redondo-Nieto, M., Arroyo, F., and Castellanos, J. (2011) Biocircuit design through engineering bacterial logic gates, Natural Computing 10, 119–127.

[32] Kylilis, N., Tuza, Z. A., Stan, G.-B., and Polizzi, K. M. (2018) Tools for engineering coordinated system behaviour in synthetic microbial consortia, Nature communications 9, 1–9.

[33] Gorochowski, T. E., Hauert, S., Kreft, J.-U., Marucci, L., Stillman, N. R., Tang, T.-Y. D., Bandiera, L., Bartoli, V., Dixon, D. O., and Fedorec, A. J. (2020) Toward engineering biosystems with emergent collective functions, Frontiers in Bioengineering and Biotechnology 8, 705.

[34] Kittleson, J. T., Wu, G. C., and Anderson, J. C. (2012) Successes and failures in modular genetic engineering, Current opinion in chemical biology 16, 329–336.

[35] Moon, T. S., Lou, C., Tamsir, A., Stanton, B. C., and Voigt, C. A. (2012) Genetic programs constructed from layered logic gates in single cells, Nature 491, 249–253.

[36] Andrianantoandro, E., Basu, S., Karig, D. K., and Weiss, R. (2006) Synthetic biology: new engineering rules for an emerging discipline, Molecular systems biology 2.

[37] Darlington, A. P., Kim, J., Jiménez, J. I., and Bates, D. G. (2018) Dynamic allocation of orthogonal ribosomes facilitates uncoupling of co-expressed genes, Nature communications 9, 1–12.

[38] Nikolados, E.-M., Weiße, A. Y., Ceroni, F., and Oyarzún, D. A. (2019) Growth defects and loss-of-function in synthetic gene circuits, ACS synthetic biology 8, 1231–1240.

[39] Goñi-Moreno, A. n., Benedetti, I., Kim, J., and de Lorenzo, V. (2017) Deconvolution of gene expression noise into spatial dynamics of transcription factor–promoter interplay, ACS synthetic biology 6, 1359–1369.

[40] Stoof, R., Wood, A., and Goni-Moreno, A. (2019) A Model for the Spatiotemporal Design of Gene Regulatory Circuits, ACS synthetic biology 8, 2007–2016.

[41] Goñi-Moreno, A., and Nikel, P. I. (2019) High-performance biocomputing in synthetic biology–integrated transcriptional and metabolic circuits, Frontiers in bioengineering and biotechnology 7.

[42] Oyarzún, D. A., and Stan, G.-B. V. (2013) Synthetic gene circuits for metabolic control: design trade-offs and constraints, Journal of The Royal Society Interface 10, 20120671.

[43] Couto, J. M., McGarrity, A., Russell, J., and Sloan, W. T. (2018) The effect of metabolic stress on genome stability of a synthetic biology chassis Escherichia coli K12 strain, Microbial cell factories 17, 8.

[44] de Lorenzo, V. (2011) Beware of metaphors: chasses and orthogonality in synthetic biology, Bioengineered bugs 2, 3–7.

[45] Danchin, A. (2008) Bacteria as computers making computers, FEMS microbiology reviews 33, 3–26.

[46] Macía, J., Posas, F., and Solé, R. V. (2012) Distributed computation: the new wave of synthetic biology devices, Trends in biotechnology 30, 342–349.

[47] Macia, J., Manzoni, R., Conde, N., Urrios, A., de Nadal, E., Solé, R., and Posas, F. (2016) Implementation of complex biological logic circuits using spatially distributed multicellular consortia, PLoS computational biology 12, e1004685.

[48] Virtanen, P., Gommers, R., Oliphant, T. E., Haberland, M., Reddy, T., Cournapeau, D., Burovski, E., Peterson, P., Weckesser, W., Bright, J., van der Walt, S. J., Brett, M., Wilson, J., Millman, K. J., Mayorov, N., Nelson, A. R. J., Jones, E., Kern, R., Larson, E., Carey, C. J., Polat, I., Feng, Y., Moore, E. W., VanderPlas, J., Laxalde, D., Perktold, J., Cimrman, R., Henriksen, I., Quintero, E. A., Harris, C. R., Archibald, A. M., Ribeiro, A. H., Pedregosa, F., van Mulbregt, P., and SciPy, C. (2020) SciPy 1.0: fundamental algorithms for scientific computing in Python, Nat Methods 17, 261–272.

[49] Fréchet, M. M. (1906) Sur quelques points du calcul fonctionnel, Rendiconti del Circolo Matematico di Palermo (1884-1940) 22, 1–72.

[50] Jekel, C. F., Venter, G., Venter, M. P., Stander, N., and Haftka, R. T. (2019) Similarity measures for identifying material parameters from hysteresis loops using inverse analysis, International Journal of Material Forming 12, 355–378.

