## Supplementary Information for "Contextual dependencies expand the re-usability of genetic inverters"

#### Reconfigurable genetic inverters using contextual dependencies

May 11, 2020

[1] Systems Biology Department, Centro Nacional de Biotecnologia-CSIC, Campus de Cantoblanco, Madrid 28049, Spain
[2] School of Computing, Newcastle University, Newcastle Upon Tyne NE4 5TG, UK
[3] Centro de Biotecnología y Genómica de Plantas, Universidad Politécnica de Madrid, Instituto Nacional de Investigación y Tecnología Agraria y Alimentaria, Pozuelo de Alarcón, 28223 Madrid, Spain

#### This supplementary information includes:

- Additional figures relating to the study.
- Additional tables relating to the study.
- Description of the codes and the data relating to the study.

### Contents

**1 Decomposing fluorescence and scattering** **3**

**2 Compatibility tables for individual strains** **3**

**3 List of constructs** **5**

**4 Context Similarity** **12**

**5 Compatibility Scoring** **16**

**6 Using the provided codes and data** **21**

### 1 Decomposing fluorescence and scattering

In our experiments we aim to pick cells in the log-phase, this causes the cell-size distribution to be constant [1]. We do however see that there is some experiment to experiment variation of the cell size distribution. To negate the influence of the cell size variation into the fluorescence measures we decompose the fluorescence values in two components: a part that is experiment-dependent and a part that is scattering dependent. We evaluate the part that is scattering dependent on the context average. The raw flow cytometry files, along with the decomposed values are available at DOI:10.25405/data.ncl.12073479.

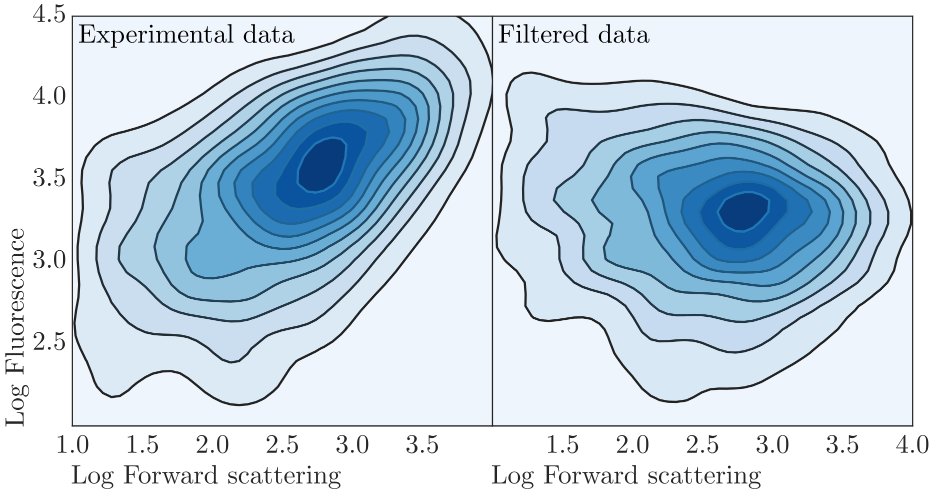

Figure S1: Removal of correlation between cell scattering and fluorescence

### 2 Compatibility tables for individual strains

In order that the genetic circuits may be treated as logic NOT gates, the continuous variable (experimentally obtained standardised fluorescence) must be interpreted as a discrete variable (representing logic 1 or 0). Thresholds are therefore required to partition the input and output fluorescence values into groups that are to be interpreted as a logic value, or rejected as ambiguous. Compatibility between two gates is a qualitative measure of the agreement of these thresholds. In particular, the output thresholds of the 'input gate' must not lie in the group of inputs that would be rejected as ambiguous by the 'output gate'. Smaller ambiguous regions will increase the numbers of compatible pairs *in silico*, but circuits built from such pairs may behave unpredictably in the presence of noise or measurement error of a real system. Larger ambiguous regions will guard against the effect of noise, at the cost of flexibility in design. In this study we use a thresholding scheme that has used previously for the same library [2]. Further, we consider that two gates may only be connected if they are compatible, and a library with many compatible gates is desirable.

Here we show the compatibility between pairs of gates in individual strains, illustrating what can be achieved by incorporating backbone as a design parameter, without changing host. The superior performance found in the DH5$\alpha$ and CC118$\lambda$pir *E. coli.* strains, in comparison to the *P.Putida* strain KT2440, may reflect the fact that the library components were initially selected for use in an *E. coli.* host, and that choice of host significantly impacts behaviour of genetic parts.

(a) Compatibility table for gates characterised in KT2440

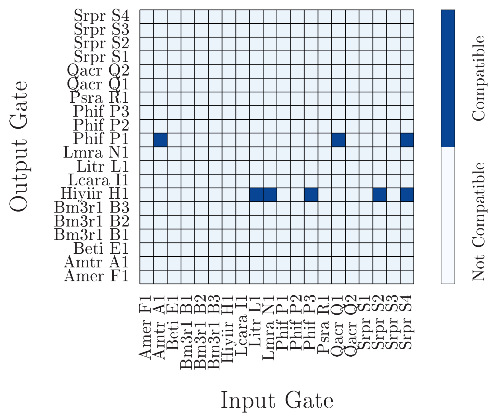

(b) Compatibility table for gates characterised in CC118λpir

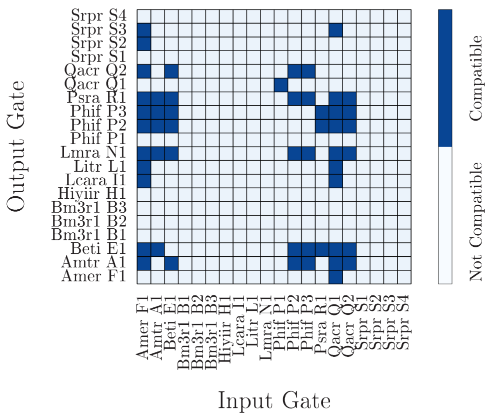

(c) Compatibility table for gates characterised in DH5α

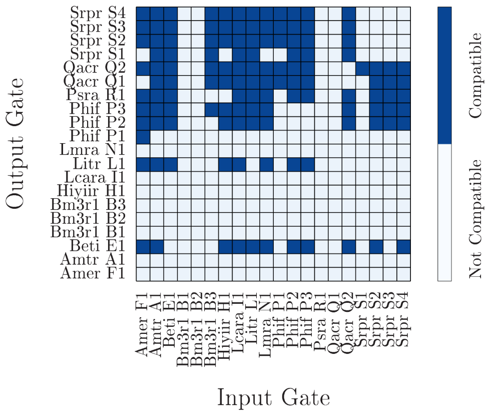

Figure S2: Compatibility tables for the library in different hosts. Input gate on the x-axis is the first gate, whose output provides the input for 'Output gate' on the y-axis. Two gates are compatible if their thresholds agree, and they do not use the same repressor molecule.

### 3 List of constructs

Table S1: 12 main gates and 8 variants that were obtained from the Cello work.

| Inverters | Description | Study |
|---|---|---|
| pAN::AmeR-F1 | NOT Gate | Nielsen et al[1] |
| pAN::AmtR-A1 | NOT Gate | Nielsen et al[1] |
| pAN::BetI-E1 | NOT Gate | Nielsen et al[1] |
| pAN::BM3R1-B1 | NOT Gate | Nielsen et al[1] |
| pAN::BM3R1-B2 | NOT Gate | Nielsen et al[1] |
| pAN::BM3R1-B3 | NOT Gate | Nielsen et al[1] |
| pAN::HIyIIR-H1 | NOT Gate | Nielsen et al[1] |
| pAN::lcaRA-I1 | NOT Gate | Nielsen et al[1] |
| pAN::LitR-L1 | NOT Gate | Nielsen et al[1] |
| pAN::LmrA-N1 | NOT Gate | Nielsen et al[1] |
| pAN::PhIF-P1 | NOT Gate | Nielsen et al[1] |
| pAN::PhIF-P2 | NOT Gate | Nielsen et al[1] |
| pAN::PhIF-P3 | NOT Gate | Nielsen et al[1] |
| pAN::PsrA-R1 | NOT Gate | Nielsen et al[1] |
| pAN::QacR-Q1 | NOT Gate | Nielsen et al[1] |
| pAN::QacR-Q2 | NOT Gate | Nielsen et al[1] |
| pAN::SrpR-S1 | NOT Gate | Nielsen et al[1] |
| pAN::SrpR-S2 | NOT Gate | Nielsen et al[1] |
| pAN::SrpR-S3 | NOT Gate | Nielsen et al[1] |
| pAN::SrpR-S4 | NOT Gate | Nielsen et al[1] |
| pAN::1201 | Autofluorescence | Nielsen et al[1] |
| pAN::1717 | Standardisation | Nielsen et al[1] |
| pAN::1818 | Promoter Activity | Nielsen et al[1] |

Table S2: List of all libraries for context dependent inverters.

| Strain | Backbone | Stock Name | Library | Work | Definition |
|---|---|---|---|---|---|
| Ecoli DH5$\alpha$ | pAN | Tas74 | Ecoli DH5$\alpha$ pAN::Amer-F1 | This study | NOT Gate |
| | | Tas75 | Ecoli DH5$\alpha$ pAN::AmtR-A1 | This study | NOT Gate |
| | | Tas76 | Ecoli DH5$\alpha$ pAN::BetI-E1 | This study | NOT Gate |
| | | Tas77 | Ecoli DH5$\alpha$ pAN::BM3R1-B1 | This study | NOT Gate |
| | | Tas78 | Ecoli DH5$\alpha$ pAN::BM3R1-B2 | This study | NOT Gate |
| | | Tas79 | Ecoli DH5$\alpha$ pAN::BM3R1-B3 | This study | NOT Gate |
| | | Tas364 | Ecoli DH5$\alpha$ pAN::HIyIIR-H1 | This study | NOT Gate |
| | | Tas81 | Ecoli DH5$\alpha$ pAN::lcaRA-I1 | This study | NOT Gate |
| | | Tas82 | Ecoli DH5$\alpha$ pAN::LitR-L1 | This study | NOT Gate |
| | | Tas83 | Ecoli DH5$\alpha$ pAN::LmrA-N1 | This study | NOT Gate |
| | | Tas84 | Ecoli DH5$\alpha$ pAN::PhIF-P1 | This study | NOT Gate |
| | | Tas85 | Ecoli DH5$\alpha$ pAN::PhIF-P2 | This study | NOT Gate |
| | | Tas86 | Ecoli DH5$\alpha$ pAN::PhIF-P3 | This study | NOT Gate |
| | | Tas87 | Ecoli DH5$\alpha$ pAN::PsrA-R1 | This study | NOT Gate |
| | | Tas88 | Ecoli DH5$\alpha$ pAN::QacR-Q1 | This study | NOT Gate |
| | | Tas89 | Ecoli DH5$\alpha$ pAN::QacR-Q2 | This study | NOT Gate |
| | | Tas90 | Ecoli DH5$\alpha$ pAN::SrpR-S1 | This study | NOT Gate |
| | | Tas91 | Ecoli DH5$\alpha$ pAN::SrpR-S2 | This study | NOT Gate |
| | | Tas365 | Ecoli DH5$\alpha$ pAN::SrpR-S3 | This study | NOT Gate |
| | | Tas93 | Ecoli DH5$\alpha$ pAN::SrpR-S4 | This study | NOT Gate |
| | | Tas94 | Ecoli DH5$\alpha$ pAN::1201 | This study | Empty plasmid for autofluorescence |
| | | Tas95 | Ecoli DH5$\alpha$ pAN::1717 | This study | RPU standard plasmid for standardization |
| | | Tas366 | Ecoli DH5$\alpha$ pAN::1818 | This study | pTac activity plasmid for promoter activity |
| | pSEVA221 | Tas385 | Ecoli DH5$\alpha$ pSeva221::Amer-F1 | This study | NOT Gate |
| | | Tas386 | Ecoli DH5$\alpha$ pSeva221::AmtR-A1 | This study | NOT Gate |
| | | Tas387 | Ecoli DH5$\alpha$ pSeva221::BetI-E1 | This study | NOT Gate |
| | | Tas388 | Ecoli DH5$\alpha$ pSeva221::BM3R1-B1 | This study | NOT Gate |
| | | Tas389 | Ecoli DH5$\alpha$ pSeva221::BM3R1-B2 | This study | NOT Gate |
| | | Tas390 | Ecoli DH5$\alpha$ pSeva221::BM3R1-B3 | This study | NOT Gate |
| | | Tas391 | Ecoli DH5$\alpha$ pSeva221::HIyIIR-H1 | This study | NOT Gate |
| | | Tas392 | Ecoli DH5$\alpha$ pSeva221::lcaRA-I1 | This study | NOT Gate |
| | | Tas393 | Ecoli DH5$\alpha$ pSeva221::LitR-L1 | This study | NOT Gate |
| | | Tas394 | Ecoli DH5$\alpha$ pSeva221::LmrA-N1 | This study | NOT Gate |
| | | Tas395 | Ecoli DH5$\alpha$ pSeva221::PhIF-P1 | This study | NOT Gate |
| | | Tas396 | Ecoli DH5$\alpha$ pSeva221::PhIF-P2 | This study | NOT Gate |
| | | Tas397 | Ecoli DH5$\alpha$ pSeva221::PhIF-P3 | This study | NOT Gate |
| | | Tas398 | Ecoli DH5$\alpha$ pSeva221::PsrA-R1 | This study | NOT Gate |
| | | Tas399 | Ecoli DH5$\alpha$ pSeva221::QacR-Q1 | This study | NOT Gate |
| | | Tas400 | Ecoli DH5$\alpha$ pSeva221::QacR-Q2 | This study | NOT Gate |
| | | Tas401 | Ecoli DH5$\alpha$ pSeva221::SrpR-S1 | This study | NOT Gate |
| | | Tas402 | Ecoli DH5$\alpha$ pSeva221::SrpR-S2 | This study | NOT Gate |
| | | Tas403 | Ecoli DH5$\alpha$ pSeva221::SrpR-S3 | This study | NOT Gate |
| | | Tas404 | Ecoli DH5$\alpha$ pSeva221::SrpR-S4 | This study | NOT Gate |

| | | | | | |
|---|---|---|---|---|---|
| | | Tas361 | Ecoli DH5$\alpha$ pSeva221::1201 | This study | Empty plasmid for autofluorescence |
| | | Tas362 | Ecoli DH5$\alpha$ pSeva221::1717 | This study | RPU standard plasmid for standardization |
| | | Tas363 | Ecoli DH5$\alpha$ pSeva221::1818 | This study | pTac activity plasmid for promoter activity |
| Ecoli cc118$\lambda$pir | pSEVA221 | Tas1 | Ecoli cc118$\lambda$pir pSeva221::Amer-F1 | This study | NOT Gate |
| | | Tas2 | Ecoli cc118$\lambda$pir pSeva221::AmtR-A1 | This study | NOT Gate |
| | | Tas3 | Ecoli cc118$\lambda$pir pSeva221::BetI-E1 | This study | NOT Gate |
| | | Tas4 | Ecoli cc118$\lambda$pir pSeva221::BM3R1-B1 | This study | NOT Gate |
| | | Tas5 | Ecoli cc118$\lambda$pir pSeva221::BM3R1-B2 | This study | NOT Gate |
| | | Tas6 | Ecoli cc118$\lambda$pir pSeva221::BM3R1-B3 | This study | NOT Gate |
| | | Tas7 | Ecoli cc118$\lambda$pir pSeva221::HIyIIR-H1 | This study | NOT Gate |
| | | Tas8 | Ecoli cc118$\lambda$pir pSeva221::lcaRA-I1 | This study | NOT Gate |
| | | Tas9 | Ecoli cc118$\lambda$pir pSeva221::LitR-L1 | This study | NOT Gate |
| | | Tas10 | Ecoli cc118$\lambda$pir pSeva221::LmrA-N1 | This study | NOT Gate |
| | | Tas11 | Ecoli cc118$\lambda$pir pSeva221::PhIF-P1 | This study | NOT Gate |
| | | Tas12 | Ecoli cc118$\lambda$pir pSeva221::PhIF-P2 | This study | NOT Gate |
| | | Tas13 | Ecoli cc118$\lambda$pir pSeva221::PhIF-P3 | This study | NOT Gate |
| | | Tas14 | Ecoli cc118$\lambda$pir pSeva221::PsrA-R1 | This study | NOT Gate |
| | | Tas15 | Ecoli cc118$\lambda$pir pSeva221::QacR-Q1 | This study | NOT Gate |
| | | Tas16 | Ecoli cc118$\lambda$pir pSeva221::QacR-Q2 | This study | NOT Gate |
| | | Tas17 | Ecoli cc118$\lambda$pir pSeva221::SrpR-S1 | This study | NOT Gate |
| | | Tas18 | Ecoli cc118$\lambda$pir pSeva221::SrpR-S2 | This study | NOT Gate |
| | | Tas19 | Ecoli cc118$\lambda$pir pSeva221::SrpR-S3 | This study | NOT Gate |
| | | Tas20 | Ecoli cc118$\lambda$pir pSeva221::SrpR-S4 | This study | NOT Gate |
| | | Tas65 | Ecoli cc118$\lambda$pir pSeva221::1201 | This study | Empty plasmid for autofluorescence |
| | | Tas68 | Ecoli cc118$\lambda$pir pSeva221::1717 | This study | RPU standard plasmid for standardization |
| | | Tas317 | Ecoli cc118$\lambda$pir pSeva221::1818 | This study | pTac activity plasmid for promoter activity |
| | pSEVA231 | Tas367 | Ecoli cc118$\lambda$pir pSeva231::Amer-F1 | This study | NOT Gate |
| | | Tas368 | Ecoli cc118$\lambda$pir pSeva231::AmtR-A1 | This study | NOT Gate |
| | | Tas23 | Ecoli cc118$\lambda$pir pSeva231::BetI-E1 | This study | NOT Gate |
| | | Tas24 | Ecoli cc118$\lambda$pir pSeva231::BM3R1-B1 | This study | NOT Gate |
| | | Tas25 | Ecoli cc118$\lambda$pir pSeva231::BM3R1-B2 | This study | NOT Gate |
| | | Tas26 | Ecoli cc118$\lambda$pir pSeva231::BM3R1-B3 | This study | NOT Gate |
| | | Tas27 | Ecoli cc118$\lambda$pir pSeva231::HIyIIR-H1 | This study | NOT Gate |
| | | Tas28 | Ecoli cc118$\lambda$pir pSeva231::lcaRA-I1 | This study | NOT Gate |
| | | Tas29 | Ecoli cc118$\lambda$pir pSeva231::LitR-L1 | This study | NOT Gate |
| | | Tas30 | Ecoli cc118$\lambda$pir pSeva231::LmrA-N1 | This study | NOT Gate |
| | | Tas31 | Ecoli cc118$\lambda$pir pSeva231::PhIF-P1 | This study | NOT Gate |
| | | Tas32 | Ecoli cc118$\lambda$pir pSeva231::PhIF-P2 | This study | NOT Gate |
| | | Tas33 | Ecoli cc118$\lambda$pir pSeva231::PhIF-P3 | This study | NOT Gate |
| | | Tas34 | Ecoli cc118$\lambda$pir pSeva231::PsrA-R1 | This study | NOT Gate |
| | | Tas369 | Ecoli cc118$\lambda$pir pSeva231::QacR-Q1 | This study | NOT Gate |
| | | Tas36 | Ecoli cc118$\lambda$pir pSeva231::QacR-Q2 | This study | NOT Gate |
| | | Tas37 | Ecoli cc118$\lambda$pir pSeva231::SrpR-S1 | This study | NOT Gate |
| | | Tas38 | Ecoli cc118$\lambda$pir pSeva231::SrpR-S2 | This study | NOT Gate |
| | | Tas39 | Ecoli cc118$\lambda$pir pSeva231::SrpR-S3 | This study | NOT Gate |
| | | Tas40 | Ecoli cc118$\lambda$pir pSeva231::SrpR-S4 | This study | NOT Gate |

| | | | | | |
|---|---|---|---|---|---|
| | | Tas66 | Ecoli cc118λpir pSeva231::1201 | This study | Empty plasmid for autofluorescence |
| | | Tas69 | Ecoli cc118λpir pSeva231::1717 | This study | RPU standard plasmid for standardization |
| | | Tas370 | Ecoli cc118λpir pSeva231::1818 | This study | pTac activity plasmid for promoter activity |
| Pputida KT2440 | pSEVA221 | Tas193 | Pputida KT2440 pSeva221::Amer-F1 | This study | NOT Gate |
| | | Tas194 | Pputida KT2440 pSeva221::AmtR-A1 | This study | NOT Gate |
| | | Tas195 | Pputida KT2440 pSeva221::BetI-E1 | This study | NOT Gate |
| | | Tas196 | Pputida KT2440 pSeva221::BM3R1-B1 | This study | NOT Gate |
| | | Tas197 | Pputida KT2440 pSeva221::BM3R1-B2 | This study | NOT Gate |
| | | Tas198 | Pputida KT2440 pSeva221::BM3R1-B3 | This study | NOT Gate |
| | | Tas199 | Pputida KT2440 pSeva221::HIyIIR-H1 | This study | NOT Gate |
| | | Tas200 | Pputida KT2440 pSeva221::lcaRA-I1 | This study | NOT Gate |
| | | Tas201 | Pputida KT2440 pSeva221::LitR-L1 | This study | NOT Gate |
| | | Tas202 | Pputida KT2440 pSeva221::LmrA-N1 | This study | NOT Gate |
| | | Tas203 | Pputida KT2440 pSeva221::PhIF-P1 | This study | NOT Gate |
| | | Tas204 | Pputida KT2440 pSeva221::PhIF-P2 | This study | NOT Gate |
| | | Tas205 | Pputida KT2440 pSeva221::PhIF-P3 | This study | NOT Gate |
| | | Tas206 | Pputida KT2440 pSeva221::PsrA-R1 | This study | NOT Gate |
| | | Tas207 | Pputida KT2440 pSeva221::QacR-Q1 | This study | NOT Gate |
| | | Tas208 | Pputida KT2440 pSeva221::QacR-Q2 | This study | NOT Gate |
| | | Tas209 | Pputida KT2440 pSeva221::SrpR-S1 | This study | NOT Gate |
| | | Tas210 | Pputida KT2440 pSeva221::SrpR-S2 | This study | NOT Gate |
| | | Tas211 | Pputida KT2440 pSeva221::SrpR-S3 | This study | NOT Gate |
| | | Tas212 | Pputida KT2440 pSeva221::SrpR-S4 | This study | NOT Gate |
| | | Tas257 | Pputida KT2440 pSeva221::1201 | This study | Empty plasmid for autofluorescence |
| | | Tas260 | Pputida KT2440 pSeva221::1717 | This study | RPU standard plasmid for standardization |
| | | Tas266 | Pputida KT2440 pSeva221::1818 | This study | pTac activity plasmid for promoter activity |
| | pSEVA231 | Tas213 | Pputida KT2440 pSeva231::Amer-F1 | This study | NOT Gate |
| | | Tas214 | Pputida KT2440 pSeva231::AmtR-A1 | This study | NOT Gate |
| | | Tas215 | Pputida KT2440 pSeva231::BetI-E1 | This study | NOT Gate |
| | | Tas216 | Pputida KT2440 pSeva231::BM3R1-B1 | This study | NOT Gate |
| | | Tas217 | Pputida KT2440 pSeva231::BM3R1-B2 | This study | NOT Gate |
| | | Tas218 | Pputida KT2440 pSeva231::BM3R1-B3 | This study | NOT Gate |
| | | Tas219 | Pputida KT2440 pSeva231::HIyIIR-H1 | This study | NOT Gate |
| | | Tas220 | Pputida KT2440 pSeva231::lcaRA-I1 | This study | NOT Gate |
| | | Tas221 | Pputida KT2440 pSeva231::LitR-L1 | This study | NOT Gate |
| | | Tas222 | Pputida KT2440 pSeva231::LmrA-N1 | This study | NOT Gate |
| | | Tas223 | Pputida KT2440 pSeva231::PhIF-P1 | This study | NOT Gate |
| | | Tas224 | Pputida KT2440 pSeva231::PhIF-P2 | This study | NOT Gate |
| | | Tas225 | Pputida KT2440 pSeva231::PhIF-P3 | This study | NOT Gate |
| | | Tas226 | Pputida KT2440 pSeva231::PsrA-R1 | This study | NOT Gate |
| | | Tas227 | Pputida KT2440 pSeva231::QacR-Q1 | This study | NOT Gate |
| | | Tas228 | Pputida KT2440 pSeva231::QacR-Q2 | This study | NOT Gate |
| | | Tas229 | Pputida KT2440 pSeva231::SrpR-S1 | This study | NOT Gate |
| | | Tas230 | Pputida KT2440 pSeva231::SrpR-S2 | This study | NOT Gate |
| | | Tas231 | Pputida KT2440 pSeva231::SrpR-S3 | This study | NOT Gate |
| | | Tas232 | Pputida KT2440 pSeva231::SrpR-S4 | This study | NOT Gate |

| | | | | | |
|---|---|---|---|---|---|
| | | Tas258 | Pputida KT2440 pSeva231::1201 | This study | Empty plasmid for autofluorescence |
| | | Tas261 | Pputida KT2440 pSeva231::1717 | This study | RPU standard plasmid for standardization |
| | | Tas267 | Pputida KT2440 pSeva231::1818 | This study | pTac activity plasmid for promoter activity |
| | pSEVA251 | Tas233 | Pputida KT2440 pSeva251::Amer-F1 | This study | NOT Gate |
| | | Tas234 | Pputida KT2440 pSeva251::AmtR-A1 | This study | NOT Gate |
| | | Tas235 | Pputida KT2440 pSeva251::BetI-E1 | This study | NOT Gate |
| | | Tas237 | Pputida KT2440 pSeva251::BM3R1-B2 | This study | NOT Gate |
| | | Tas238 | Pputida KT2440 pSeva251::BM3R1-B3 | This study | NOT Gate |
| | | Tas239 | Pputida KT2440 pSeva251::HIyIIR-H1 | This study | NOT Gate |
| | | Tas240 | Pputida KT2440 pSeva251::lcaRA-I1 | This study | NOT Gate |
| | | Tas241 | Pputida KT2440 pSeva251::LitR-L1 | This study | NOT Gate |
| | | Tas242 | Pputida KT2440 pSeva251::LmrA-N1 | This study | NOT Gate |
| | | Tas243 | Pputida KT2440 pSeva251::PhIF-P1 | This study | NOT Gate |
| | | Tas244 | Pputida KT2440 pSeva251::PhIF-P2 | This study | NOT Gate |
| | | Tas246 | Pputida KT2440 pSeva251::PsrA-R1 | This study | NOT Gate |
| | | Tas247 | Pputida KT2440 pSeva251::QacR-Q1 | This study | NOT Gate |
| | | Tas249 | Pputida KT2440 pSeva251::SrpR-S1 | This study | NOT Gate |
| | | Tas252 | Pputida KT2440 pSeva251::SrpR-S4 | This study | NOT Gate |
| | | Tas259 | Pputida KT2440 pSeva251::1201 | This study | Empty plasmid for autofluorescence |
| | | Tas262 | Pputida KT2440 pSeva251::1717 | This study | RPU standard plasmid for standardization |
| | | Tas285 | Pputida KT2440 pSeva251::1818 | This study | pTac activity plasmid for promoter activity |

Table S3: Primers List

| Name | Sequence 5→3 | Notes |
|---|---|---|
| Gate-Ctrl-Unvrsl-1 | TCTAGGGCGGCGGATTTG | This is the position right before the MCS. Same for pSEVA221, pSEVA231, pSEVA251 |
| Gate-Ctrl-Unvrsl-3 | TGGTGAGCAAGGGCGAG | This is the position right after the multiplicity region of the gates This is same for pSEVA221, pSEVA231, pSEVA251 |
| Gate-Ctrl-Unvrsl-4 | ACCTTAGCTACCAGTCCGC | This is the position is same for pSEVA221, pSEVA231, pSEVA251 |
| Gate-Ctrl-Unvrsl-5 | ACAATCTTCTCGCGCAACG | This is the position is same for pSEVA221, pSEVA231, pSEVA251 |
| Gate-Ctrl-Unvrsl-6 | CGGTGAAGGGCAATCAGCT | This is the position is same for pSEVA221, pSEVA231, pSEVA251 |
| Gate-Ctrl-Unvrsl-7 | GAATGCATTATATGCACTCAGCGC | This is the position is the last one for covering the integrated region (when considered DNApol moves at least 600 bp) and same for pSEVA221, pSEVA231, pSEVA251 |
| Gate-Ctrl-2-AmeR-F1 | TTTTAGCAGCAAAAACGCACTGG | This is the 500th position for the AmeR-F1 gate |
| Gate-Ctrl-2-AmtR-A1 | CCTGCTGAAAAGCACCGTTG | This is the 500th position for the AmtR-A1 gate |
| Gate-Ctrl-2-BetI-E1 | GTCTGCATGCACTGCCG | This is the 500th position for the BetI-E1 gate |
| Gate-Ctrl-2-BM3R1-B123 | CGGTCTGGCAAATGAACGTG | This is the 500th position for the BM3R1-B1 BM3R1-B2 and BM3R1-B3 gates and can be used in every cloning with these gates |
| Gate-Ctrl-2-HIyRII-H1 | TGCCGAACTTTCTGGAAAAAAACC | This is the 500th position for the HIyIIR-H1 gate |
| Gate-Ctrl-2-lcaRA-I1 | ACAAAAGCAACTATAGCATCGATGC | This is the 500th position for the lcaRA-I1 gate |
| Gate-Ctrl-2-litR-L1 | CTGAACAAAGTGGAAAACGAGTTTCAC | This is the 500th position for the LitR-L1 gate |
| Gate-Ctrl-2-LmrA-N1 | AAGAATATATCCGCCAGAAAATCGC | This is the 500th position for the LmrA-N1 gate |
| Gate-Ctrl-2-PhIF-P123 | AAAATGAAAGCGAACAGGTGCG | This is the 500th position for the PhIF-P1 PhIF-P2 and PhIF-P3 gates |
| Gate-Ctrl-2-PsrA-R1 | GGAACGTGAACTGGAACGTC | This is the 500th position for the PsrA-R1 gate |
| Gate-Ctrl-2-QacR-Q123 | CAAATCAAATGCAAAACCAACCGC | This is the 500th position for the QacR-Q1 and QacR-Q2 gates |
| Gate-Ctrl-2-SrpR-S1234 | TCCGCTGGAACTGGATTTTACAC | This is the 500th position for the SrpR-S1 SrpR-S2 SrpR-S3 and SrpR-S4 gates |

| Name | Sequence 5→3 | Notes |
|---|---|---|
| PS1 | AGGGCGGCGGATTTGTCC | |
| PS2 | GCGGCAACCGAGCGTTC | |
| PS3 | GAACGCTCGGTTGCCGC | |
| PS4 | CCAGCCTCGCAGAGCAGG | |
| PS5 | CCCTGCTTCGGGGTCATT | |
| PS6 | GGACAAATCCGCCGCCCT | |
| Gate to pSEVA | CCTAGATTAATTAAAACACCCTTGTAT TACTGTTTATGTAAGC | Forward primer in order to pop out the gate |
| Gate to pSEVA | GTCTAAACTAGTCGTCCGGCGTAGAG GATC | Reverse primer in order to pop out the gate |

Table S3: Primers List

In Table S1 there are 23 plasmids in total that are acquired from Cello study. 20 of these plasmids are NOT gates that have functional segments in common in all constructs. Those are LacI/$P_{tac}$ expression system inducible with IPTG and *yfp* gene that is used for output readout. Gate specific units that are not shared for all are the RBS sites and gate proteins. Plasmids have kanamycin resistance gene and p15A origin of replication.

Autofluorescence plasmid is named as pAN::1201. This plasmid is used to measure background noise created by the backbone. The plasmid is composed of a constitutive $P_{lac}$ promoter expressing lacZ$\alpha$ gene. Also, two sensor proteins TetR and LacI are present at the same operon under $P_{lacI}$ promoter. RPU standard plasmid is pAN::1717. This plasmid shares the same sensory proteins as in autofluorescence plasmid with the same $P_{lacI}$/LacI/TetR expression system, but the difference is that a *yfp* gene is expressed under constitutive promoter J23101. L3S2P21 terminator is used at the downstream of *yfp* for insulation of transcriptional read-throughs and native AraC terminator is used for insulation of *tetR* gene. pAN::1818 is the promoter activity plasmid. $P_{lacI}$ constitutive promoter controls the expression of *lacI* and *tetR* genes. *yfp* gene is expressed at the downstream of $P_{tac}$ promoter that is inducible with IPTG.

Table S2 shows the complete list of strains that are used in this study. We have constructed 7 different libraries changing in plasmid backbones, copy numbers and hosts. 7 libraries are consisting of 2 *Escherichia coli* strains (DH5$\alpha$ and CC118$\lambda$pir) and 1 *Pseudomonas putida* strain (KT2440). DH5a is composed of two different backbones pAN and pSEVA. CC118$\lambda$pir, whereas, it contains only pSEVA backbones however in two different copy numbers low and medium copy. Finally, KT2440 consists of low, medium and high copy numbers in pSEVA backbones. All 7 libraries have each of 20 NOT gates and 3 additional gates, autofluorescence plasmid, RPU standardization plasmid and Promoter activity plasmid. Only in KT2440 pSEVA251 (high copy) library we have encountered cloning problems in 5 of the gates which may have been resulted from a possible toxicity effect caused by them in higher copies. These failed gate clones are in pSEVA251 backbone BM3R1-B1, PhIF-P3, QacR-Q2, SrpR-S2 and SrpR-S3. Hence, in total 135 strains are generated in this study with NOT gates, and 21 strains that are to help to standardize outcomes and measure promoter activity, in total 156 strains listed in Table S2 are used in this work.

Table S3 gives detailed information about primers used in this study. There are four group of primers used here. First group is Gate-Unvrsl-Number primers, which are targeting various preserved regions in the expression system. These primers are to use for verification purposes of the clones and they are 6 in total. Second group of primers are called as Gate-Ctrl-2-Names. There are 12 of these plasmids which are named after the gate they are targeting. PCR verification of cloned gate can be done by using its corresponding group 2 primer. Third group is named with nomenclature PS-Number and they are 6 in total. These are general SEVA control primers targeting different regions of SEVA plasmids that help to confirm successful clones. Finally, in the fourth group there is a primer set used for pop-up of gates from pAN backbone to pSEVA backbone. These primers consist of gate homologous region and a flanking part that has PacI and SpeI restriction sides used for cloning into pSEVAs.

### 4 Context Similarity

This study found that the same genetic logic gate can exhibit differing behaviour depending upon the context in which it was situated. The characterisation of the genetic logic gate may be both quantitatively and quantitatively different. Qualitative changes, such as to the shape of the response curve, are particularly interesting when considering the effects of context-circuit interplay, because they suggest these interactions are nonlinear phenomena.

We attempted to quantify changes in curve shape using a similarity measure as described in Methods of the manuscript. Log transformation of the curve ensures that deviations in the upper regions of input and output are not disproportionately penalised. The min-max normalisation in both input and output dimensions captures the shape information of the curve. Methods based on comparison of the gradient of the curves were also considered, and produce similar results.

(a) Similarity scores heatmap for Amer F1 in 7 contexts.

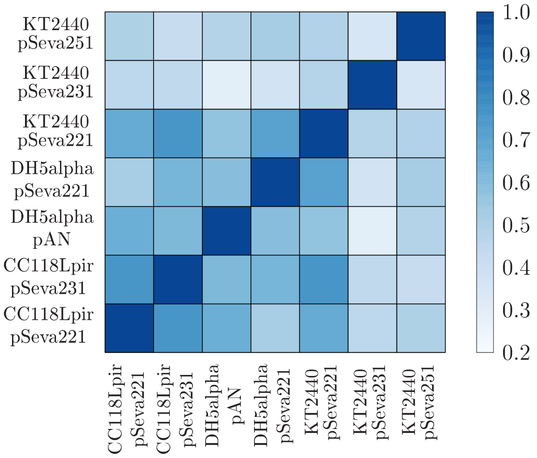

(b) Similarity scores heatmap for Amtr A1 in 7 contexts.

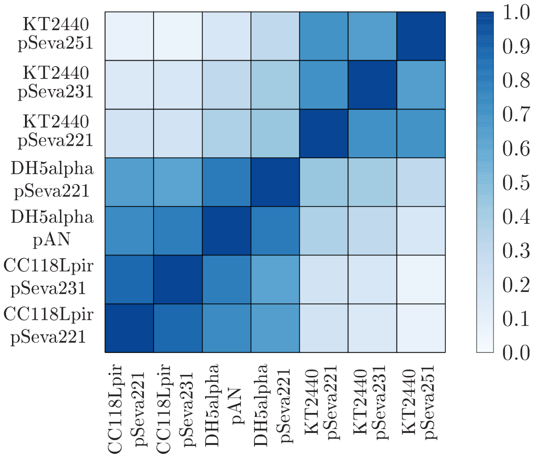

(c) Similarity scores heatmap for Beti E1 in 7 contexts.

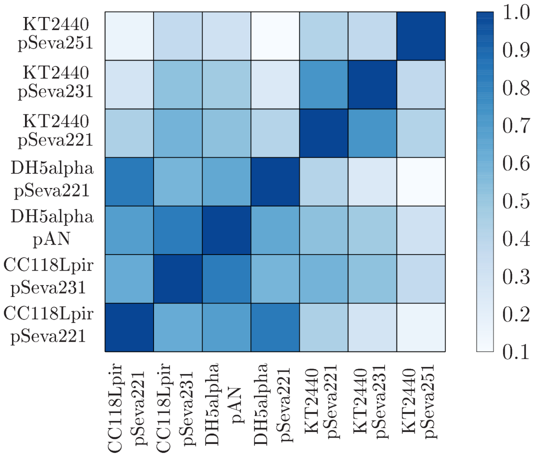

(d) Similarity scores heatmap for Bm3r1 B1 in 7 contexts.

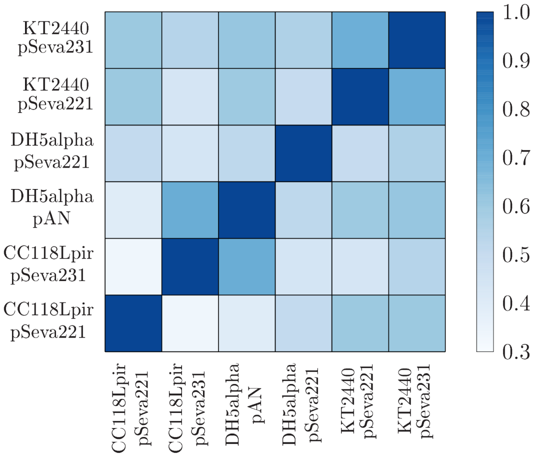

(e) Similarity scores heatmap for Bm3r1 B2 in 7 contexts.

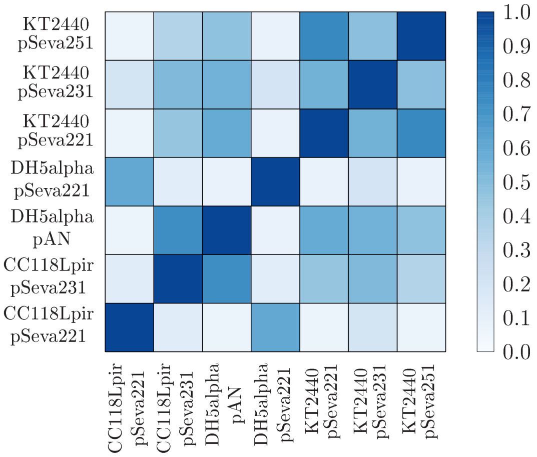

(f) Similarity scores heatmap for Bm3r1 B3 in 7 contexts.

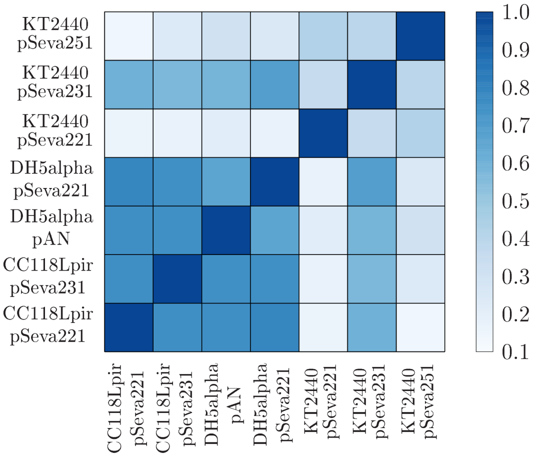

(g) Similarity scores heatmap for Hiyiir H1 in 7 contexts.

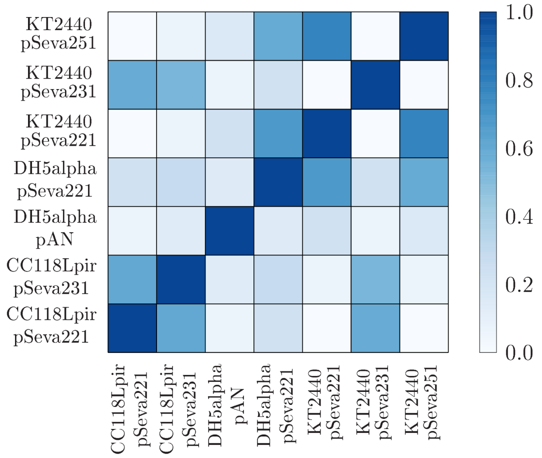

(h) Similarity scores heatmap for Lcara I1 in 7 contexts.

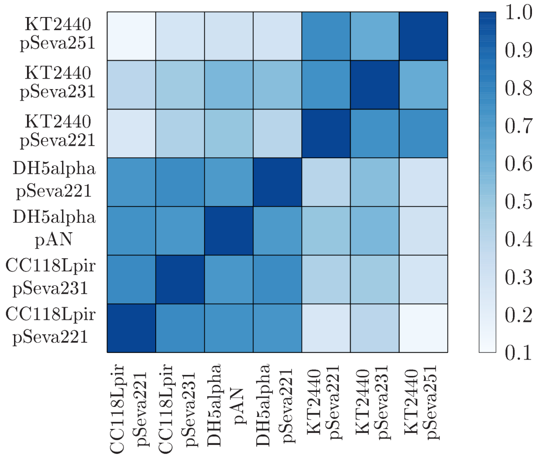

(i) Similarity scores heatmap for Litr L1 in 7 contexts.

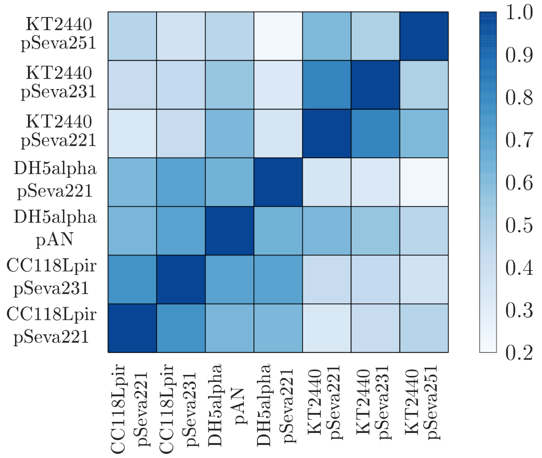

(j) Similarity scores heatmap for Lmra N1 in 7 contexts.

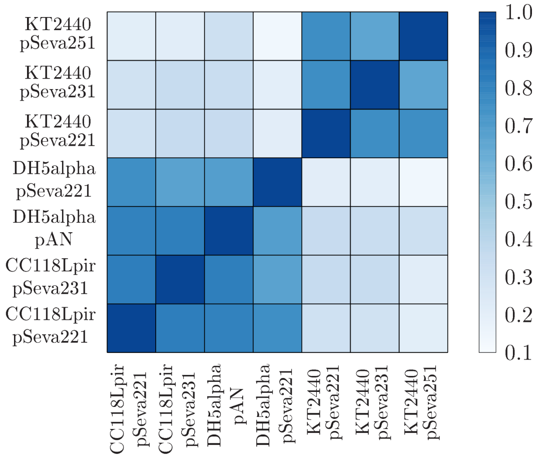

(k) Similarity scores heatmap for Phif P1 in 7 contexts.

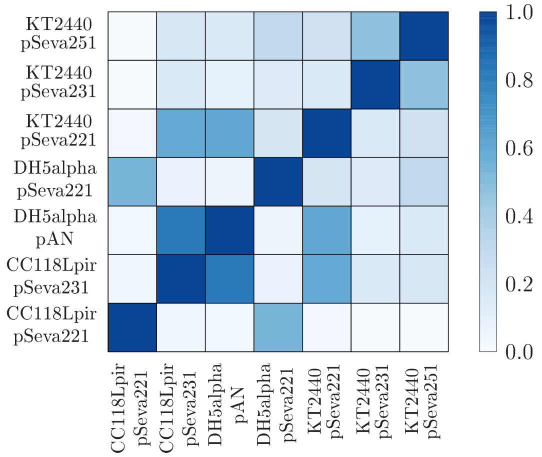

(l) Similarity scores heatmap for Phif P2 in 7 contexts.

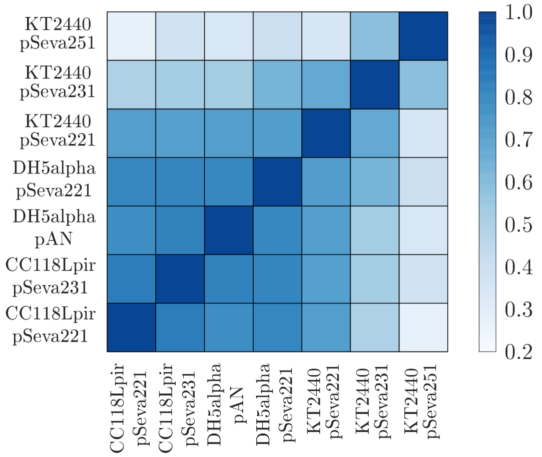

(m) Similarity scores heatmap for Phif P3 in 7 contexts.

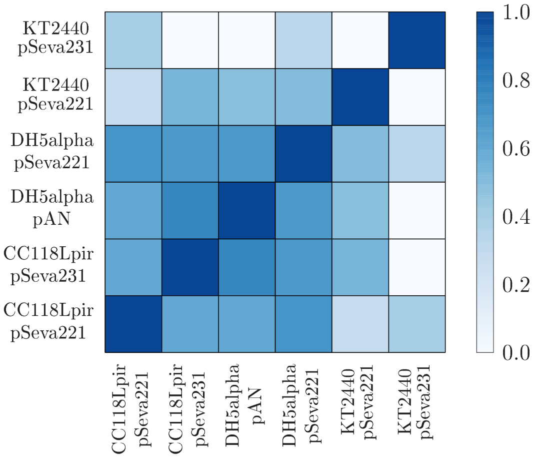

(n) Similarity scores heatmap for Psra R1 in 7 contexts.

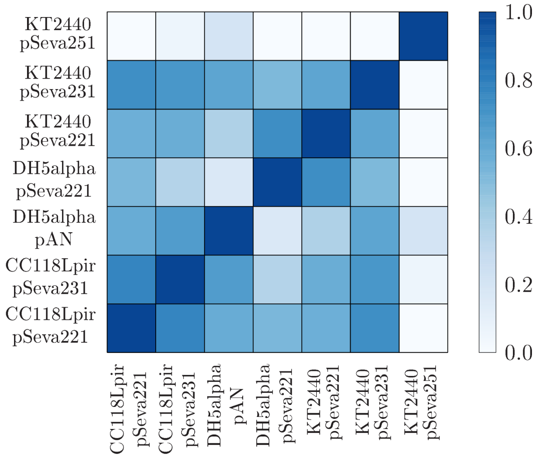

(o) Similarity scores heatmap for Qacr Q1 in 7 contexts.

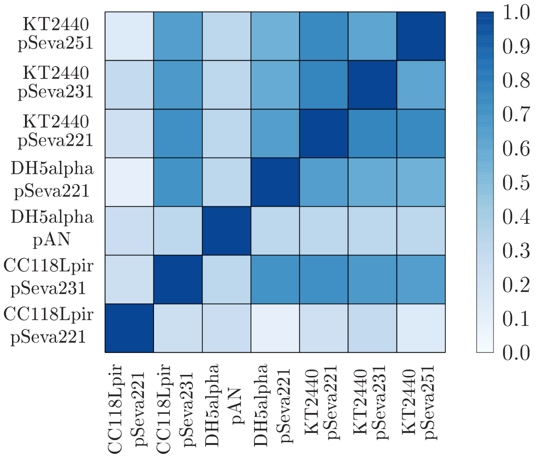

(p) Similarity scores heatmap for Qacr Q2 in 7 contexts.

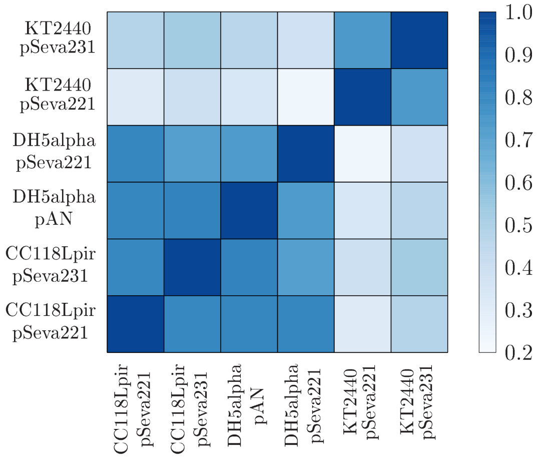

(q) Similarity scores heatmap for Srpr S1 in 7 contexts.

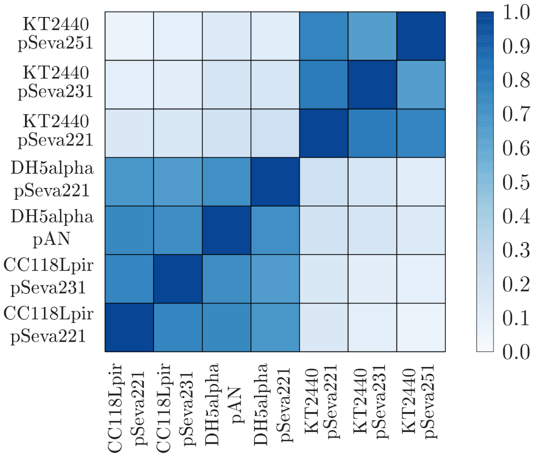

(r) Similarity scores heatmap for Srpr S2 in 7 contexts.

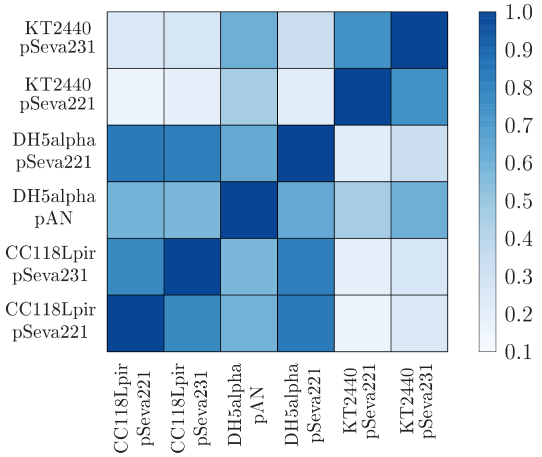

(s) Similarity scores heatmap for Srpr S3 in 7 contexts.

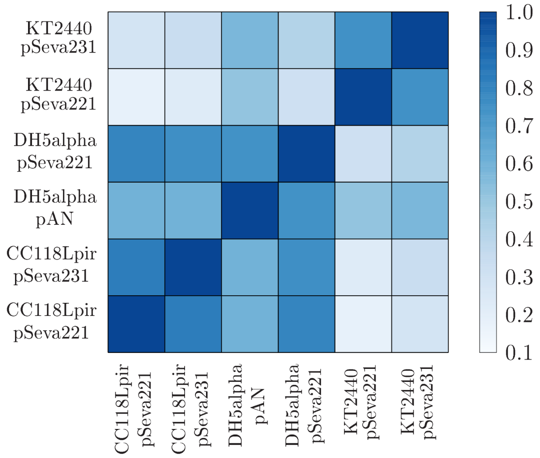

(t) Similarity scores heatmap for Srpr S4 in 7 contexts.

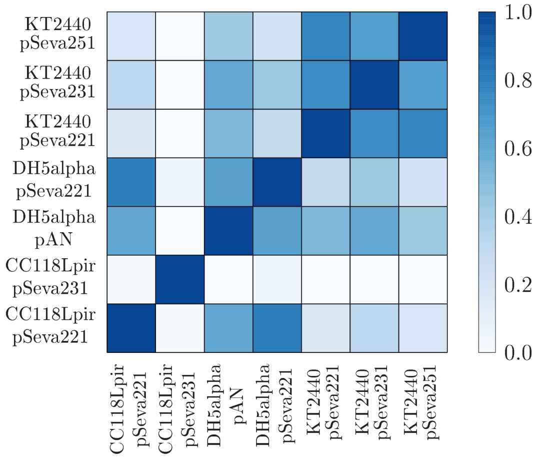

Figure S3: Similarity scores shown for all gates in all 7 contexts. A high similarity score (darker squares) is best. The minimum and maximum scores are 0 and 1, respectively.

### 5 Compatibility Scoring

Whilst compatibility tables indicate which pairs of gates may be connected, they offer no indication as to which pairs are most or least compatible. It may be desirable, from a optimisation perspective, to select pairs of gates for which the first's output thresholds lie as far away from the second's ambiguous region as possible whose. Doing so will improve performance in spite of noise and provide a greater margin for error.

Conversely, we may wish to optimise the library for a specific context by redesign of the parts. In this case, we would like to know which pairs would be compatible with only small changes to their existing behaviour, such that the reward for our optimisation efforts are maximised.

We computed a compatibility score to measure these characteristics, as defined in Methods. A positive/negative score for a pair of gates indicated the pair is compatible/incompatible. Further, a positive score represents the minimum of the maximum perturbations to the thresholds that could be tolerated, whilst retaining the compatibility of the pair. A negative score represents the maximum of the minimum perturbations to the thresholds that would be required to make the pair compatible that could be allowed to any of thresholds. Thus, pairs of gates can be ranked in terms of compatibility.

(a) Compatibility score heatmap for gates in the CC118λpir host with the pSeva221 backbone.

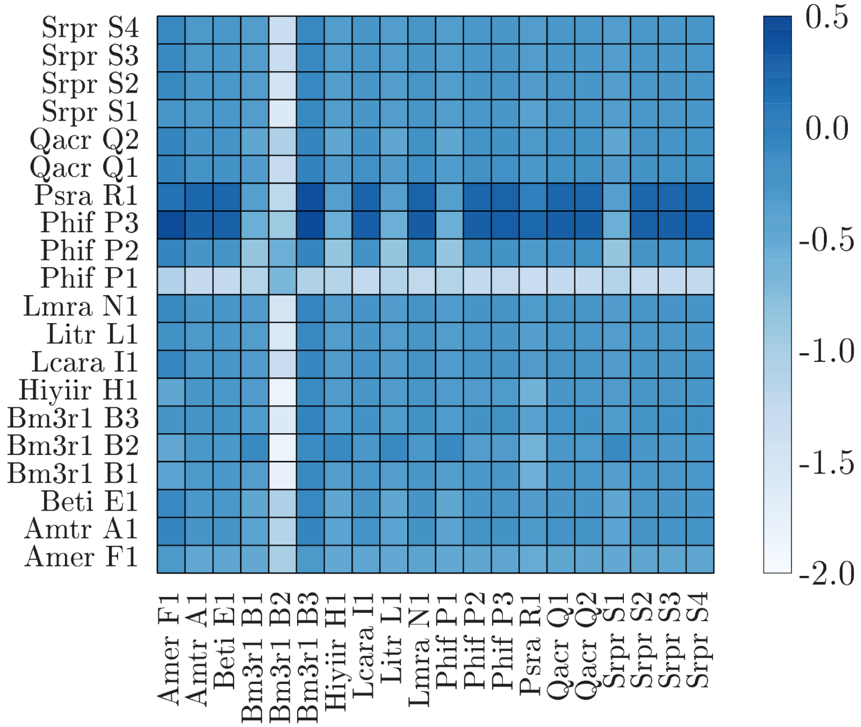

(b) Compatibility score heatmap for gates in the CC118λpir host with the pSeva231 backbone.

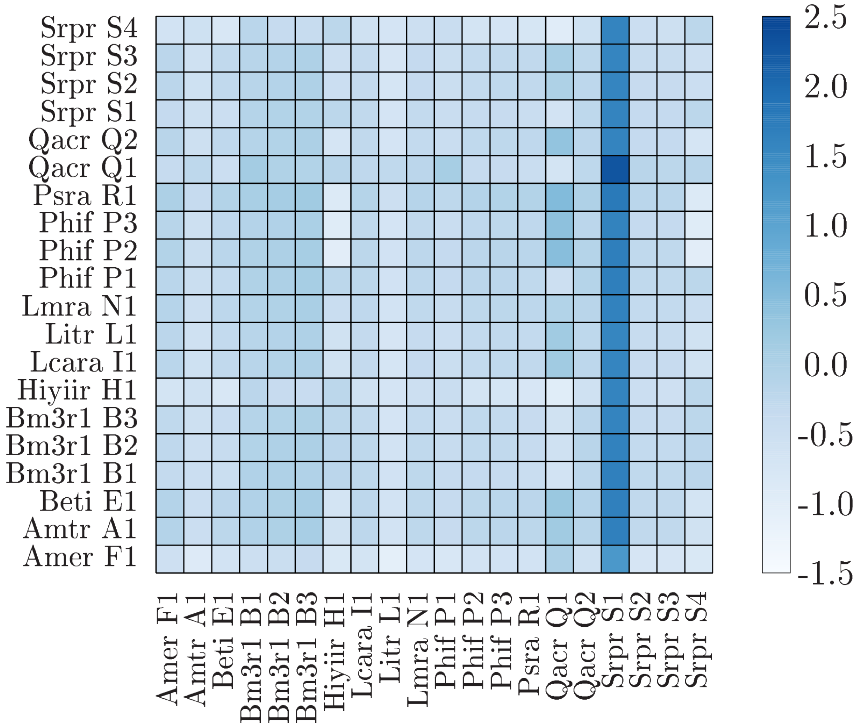

(c) Compatibility score heatmap for gates in the DH5$\alpha$ host with the pAN backbone.

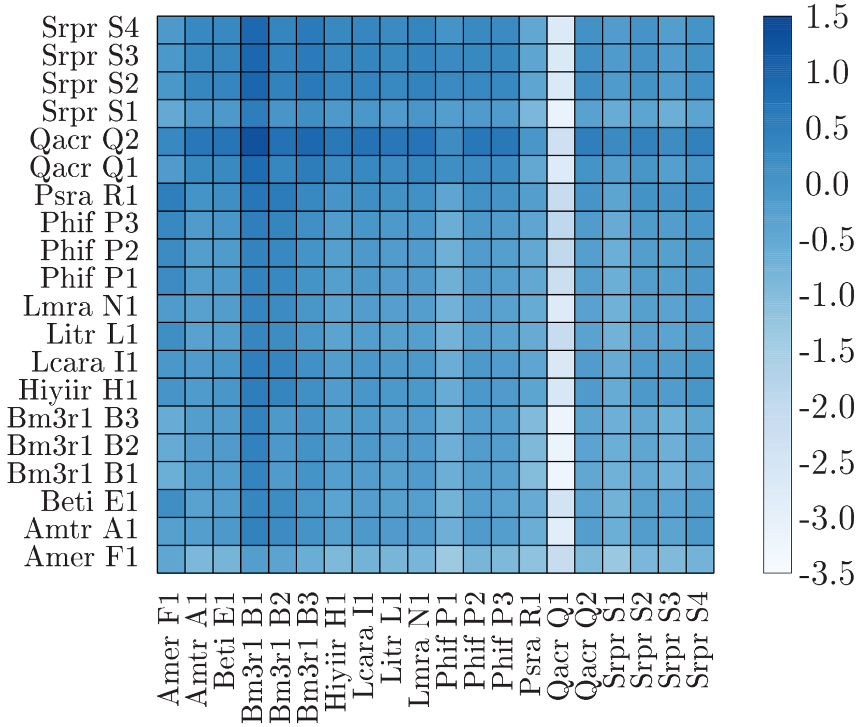

(d) Compatibility score heatmap for gates in the DH5$\alpha$ host with the pSeva221 backbone.

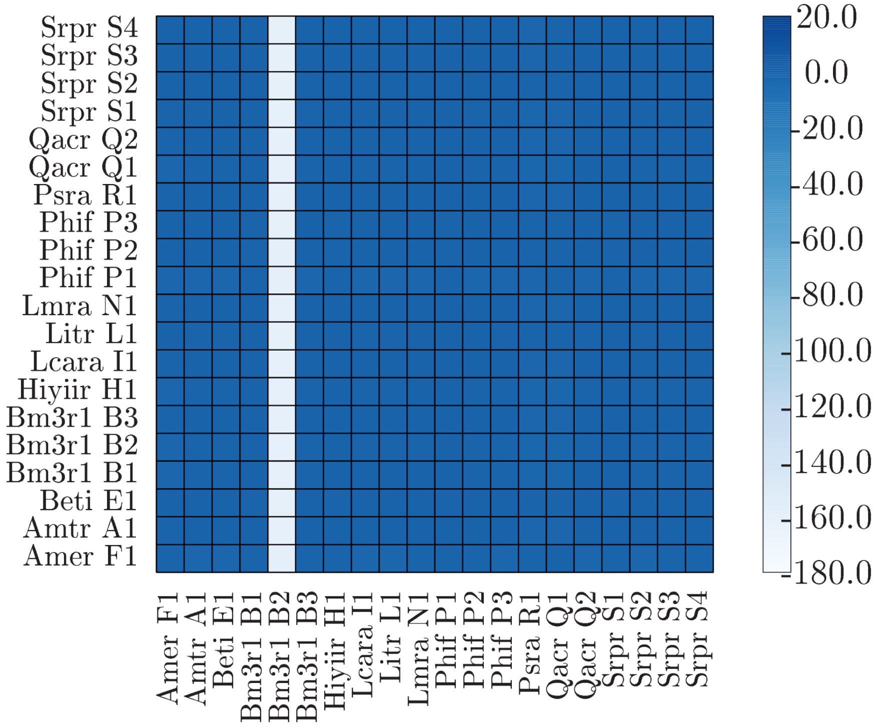

(e) Compatibility score heatmap for gates in the KT2440 host with the pSeva221 backbone.

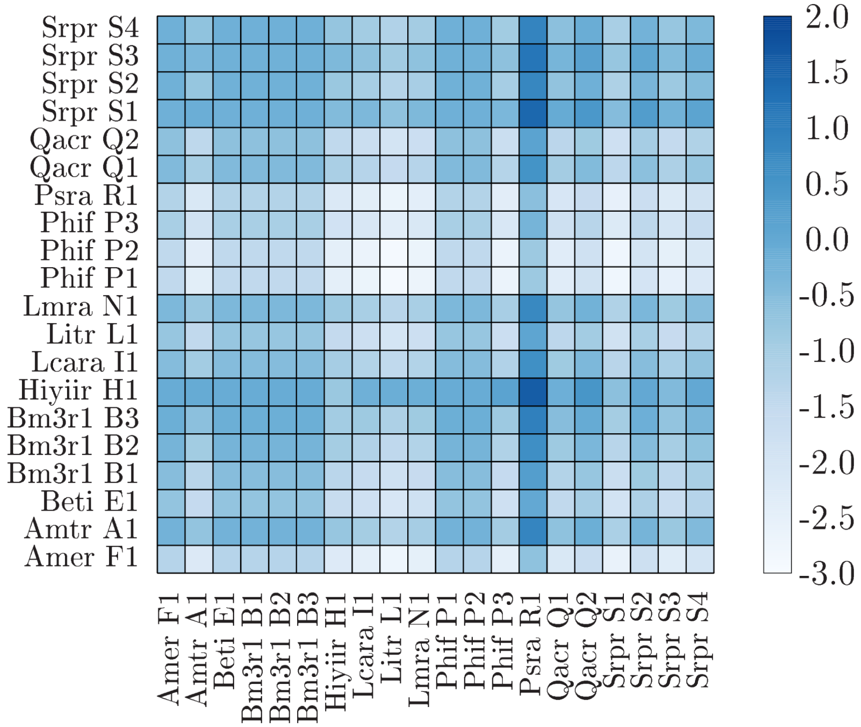

(f) Compatibility score heatmap for gates in the KT2440 host with the pSeva231 backbone.

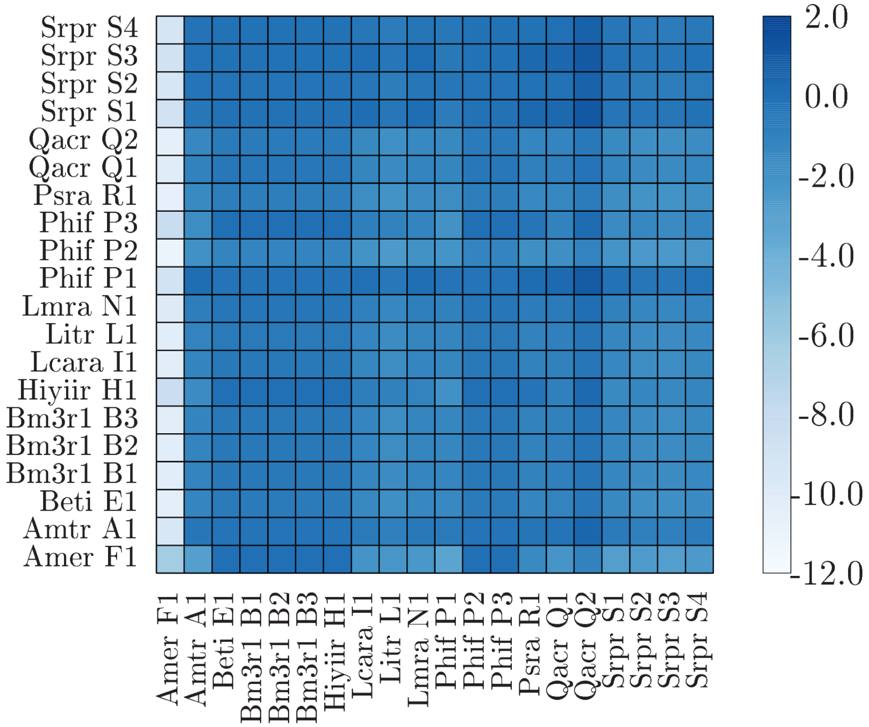

(g) Compatibility score heatmap for gates in the KT2440 host with the pSeva251 backbone.

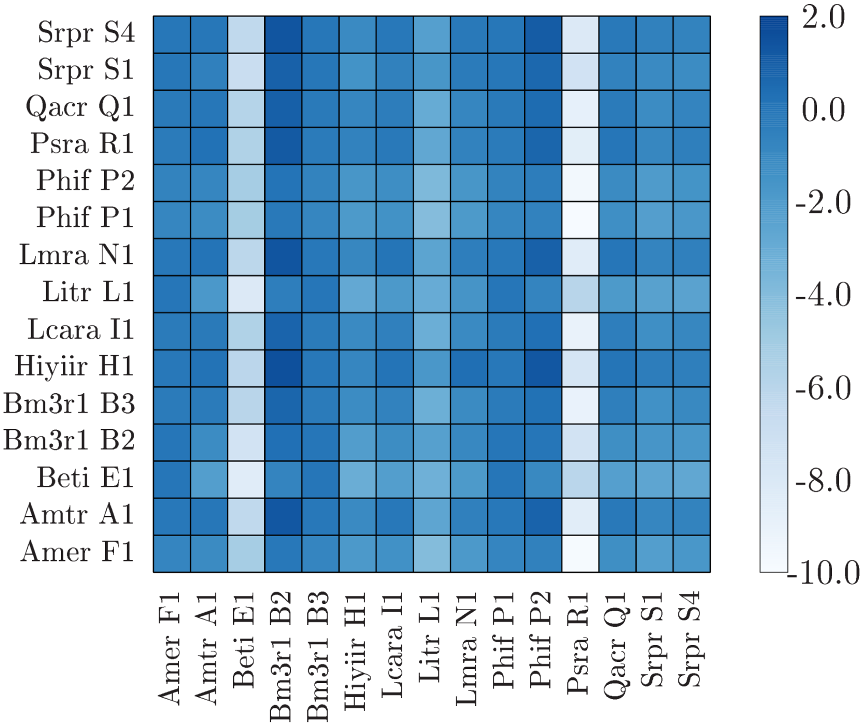

Figure S4: Compatibility scores between pairs of gates for all contexts. Higher scores are better and indicate more compatible pairs. Negative scores indicate incompatible pairs.

### 6 Using the provided codes and data

The codes that implement the *in silico* methods from this study can be found and downloaded at `https://github.com/lgrozinger/pyolin`, along with instructions detailing its use.

In order to obtain the results from this particular study, only Python3.6 or greater is needed. However, to generate the figures used in the manuscript further requires gnuplot and LaTeX. Once these dependencies are installed, the command:

```
python3 produce_figures.py full-update
```

once run from the project root directory will perform the analysis and place figures in the appropriate subdirectories. Docker users may find it more convenient to use the Dockerfile associated with the project.

It should also possible to run the same analysis on a different dataset, if the data is provided as a csv file with the expected fields and format. The processed data from this study that is provided with the codes can act as a guideline.

More information can be found in the project README at `https://github.com/lgrozinger/pyolin/blob/master/README.md`.

The data used for this study can be obtained at `https://figshare.com/s/18e6a10d708d15839837`. More information can be found in the accompanying README file.

### References

[1] P R Painter and a G Marr. Mathematics of microbial populations. *Annu. Rev. Microbiol.*, 22(January):519–548, oct 1968.

[2] Alec A. K. Nielsen, Bryan S. Der, Jonghyeon Shin, Prashant Vaidyanathan, Vanya Paralanov, Elizabeth A. Strychalski, David Ross, Douglas Densmore, and Christopher A. Voigt. Genetic circuit design automation. *Science*, 352(6281), 2016.
